# Multi-omics investigations uncover unique pathogenic markers in clinical *Klebsiella pneumoniae* that could be leveraged as novel antimicrobial targets

**DOI:** 10.1101/2023.12.22.573023

**Authors:** Lena-Sophie Swiatek, Kristin Surmann, Elias Eger, Justus Ursus Müller, Manuela Gesell Salazar, Sebastian Guenther, Guido Werner, Nils-Olaf Hübner, Jürgen A. Bohnert, Karsten Becker, Stefan E. Heiden, Uwe Völker, Michael Schwabe, Katharina Schaufler

**Affiliations:** Department of Epidemiology and Ecology of Antimicrobial Resistance, Helmholtz Institute for One Health, Helmholtz Centre for Infection Research HZI, Greifswald, Germany; Department of Functional Genomics, Interfaculty Institute for Genetics and Functional Genomics, University Medicine Greifswald, Germany; Pharmaceutical Biology, Institute of Pharmacy, University of Greifswald, Greifswald, Germany; Division 13 Nosocomial Pathogens and Antibiotic Resistances, Department of Infectious Diseases, Robert Koch Institute, Wernigerode, Germany; Central Unit for Infection Prevention and Control, University Medicine Greifswald, Greifswald, Germany.; Friedrich Loeffler-Institute of Medical Microbiology, University Medicine Greifswald, Greifswald, Germany.; University Medicine Greifswald, Greifswald, Germany

**Keywords:** Genomics, Transcriptomics, Proteomics, *Klebsiella*, Multidrug resistance, Anti-virulence strategies

## Abstract

**Background:** *Klebsiella pneumoniae* (KP), often multidrug-resistant (MDR), is a significant public health concern and frequently associated with various diseases including urinary-tract infection. In addition, in recent years, an increasing number of studies reports on the emergence of convergent KP that combine MDR with hypervirulence leading to severely limited treatment options and thus calling for alternative approaches.

**Methods:** In this study, we compared high-risk clonal KP lineages with less pathogenic *Klebsiella variicola* (KV) and *Klebsiella quasipneumoniae* (KQ) strains on multiple-omics levels and performed integrative data analysis to identify unique markers that could be subsequently leveraged as novel targets in alternative treatment strategies.

**Results:** Our initial genomic analysis revealed 107 genes as part of the patho-core genome in eight clinical KP that were associated with different metabolic pathways. Subsequent transcriptome and proteome analyses in infection-mimicking media demonstrated similar regulatory patterns among KP vs. other *Klebsiella* strains, again with metabolic responses playing a pivotal role. In total, we identified 193 KP-specific, differentially expressed genes on transcriptomic and/or proteomic levels. When then comparing these regulated genes to over 6,000 publicly available *Klebsiella* genomes, we identified unique markers either in KP genomes or adaptively regulated on transcriptomics and/or proteomics levels. An example for the latter was a gene cluster for the cellobiose phosphotransferase system that has been previously described in the context of bacterial virulence and biofilm formation.

**Conclusion:** In conclusion, our study not only highlights that KP strains demonstrate metabolic flexibility in response to particular environmental conditions, which is potentially important for their success as opportunistic pathogens, but identified unique KP-markers. Subsequent studies are needed to explore whether these markers might be prospectively used as novel anti-virulence targets, providing alternatives to traditional antibiotics.

## 1. Background

*Klebsiella* (*K*). *pneumoniae* infections pose a significant burden on public health [1], emphasized by the World Health Organization, which considers carbapenem-resistant and extended-spectrum β-lactamases (ESBL)-producing *Enterobacterales* as particularly “critical” pathogens that are often multidrug-resistant (MDR) [2]. *K. pneumoniae* colonizes human mucosal surfaces, such as the nasopharynx and gastrointestinal tract, which represent important reservoirs for subsequent infection [3]–[6]. *K. pneumoniae* is an opportunistic pathogen [7] and associated with a variety of diseases, including pneumonia, urinary tract infection (UTI), pyogenic liver abscess, meningitis, and bloodstream infections [7]–[11]. According to a recent systematic analysis of the global burden of bacterial antimicrobial resistance (AMR), MDR *K. pneumoniae* accounted for over 600,000 deaths worldwide in 2019 [1]. It is also the species responsible for most outbreaks and clinical complications within the *Klebsiella* genus [12], [13].

Traditionally, *K. pneumoniae* has been divided into two pathotypes, namely classical *K. pneumoniae* (cKp) and hypervirulent *K. pneumoniae* (hvKp) [14]. cKp strains are often resistant against several classes of antibiotics, with resistance mechanisms including mostly plasmid-mediated expression of ESBL and carbapenemases such as the New Delhi metallo-β-lactamase 1 (NDM-1), oxacillinase-48 (OXA-48), and *Klebsiella pneumoniae* carbapenemase [15]–[17], upregulation of efflux pumps [18], and mutations in membrane porins [19] or regulatory genes [20]. In contrast to the MDR characteristics of cKP, which mostly cause hospital-acquired infections, hvKp strains are susceptible to most antibiotics and frequently associated with severe infection in the healthy community [21]–[24]. Strains belonging to the hvKp pathotype often express several virulence factors, and are traditionally defined by a positive string test and thus, hypermucoviscosity [21]–[23]. However, the presence of biomarkers (i.e., *peg-344*, *iroB, iucA, rmpA*) and *in vivo* virulence is more accurate in classifying hvKp [22]. Alarmingly, an increasing number of studies report on the emergence of convergent pathotypes that combine hypervirulence with AMR features [21], [25]–[27], even against the recently approved antibiotic cefiderocol [28]. Based on multi-locus sequence typing (MLST), multiple *K. pneumoniae* sequence types (STs) have been identified, which differ in their global success and distribution [29], [30]. Among the most successful ones are international high-risk clonal lineages belonging to ST15, ST147 and ST258 mostly associated with the cKP pathotype, but, for example, also ST307, which includes converging strains as demonstrated in one of our previous studies [26], [29], [31].

Besides *K. pneumoniae* (KpI, KP), the *K. pneumoniae* species complex (KpSC) comprises other species, e.g., *K. quasipneumoniae* (KpII, KQ), and *K. variicola* (KpIII, KV) [12], [13], [32]. Despite the possibility that KQ and KV strains acquire AMR features and are marginally involved in clinical outbreaks, they are considered less pathogenic than KP, and are often described as colonizers and commensals [13], [24], [33]. For example, several studies have previously shown that the majority of KQ and KV were negative for the important virulence-associated regulator of the mucoid phenotype *rmpA* [34], [35]. The large diversity among the KpSC is reflected in a complex accessory genome that includes fitness and virulence factors related to capsule and lipopolysaccharide production, iron acquisition, and many more [13], [24]. These genes constitute great opportunities for novel target identification and utilization in alternative treatment strategies, ideally approached by multi-omics investigations [36], [37].

Overall, this highlights the urgent need for new and effective treatment options such as anti-virulence strategies, aiming to “disarm” pathogenic bacteria [36], [38]. Anti-virulence approaches exhibit several advantages compared to conventional antibiotic therapy, including the availability of a large number of targets, less evolutionary pressure, minor impact on the host commensal flora, and protection of immuno-compromised individuals [36], [38], [39]. There are several examples for successful anti-virulence applications, e.g., the inhibition of bacterial adhesion and colonization to epithelial cells with mannosides in UTI [40] or the toxin neutralization of *Clostridioides difficile* with antibodies [41].

For this study, we applied genomics, transcriptomics, and proteomics in infection-mimicking conditions. By comparing strains of international high-risk clonal KP lineages to the clinically less successful KV species, we aimed at identifying markers that are unique to pathogenic *Klebsiella* and which could be perspectively leveraged as novel targets for anti-virulence strategies.

## 2. Material and Methods

### 2.1. Bacterial strains

For our pangenome analysis, we meticulously selected eight KP STs, all characterized as MDR, representing global high-risk clonal lineages [29] (**Table 1**). Additionally, KQ and KV, recognized as commensal representatives, were included based on previous studies [13], [24]. For in-depth transcriptomics, proteomics, and phenotypic analysis, two human isolates from each of four pathogenic STs (ST15, ST147, ST258, and ST307), along with two KV strains, were selected (**Table 1**).

**Table 1:**
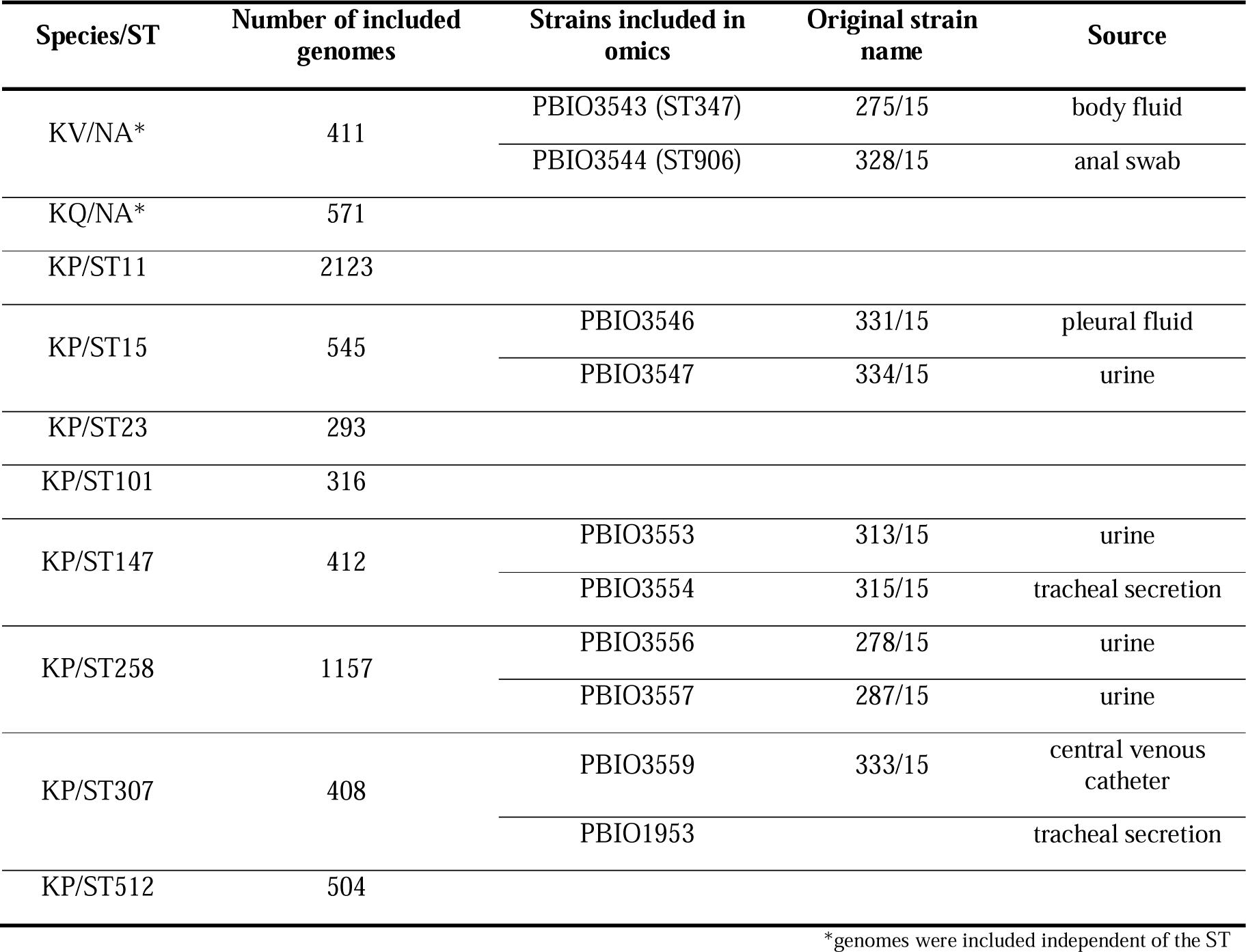
Overview of *Klebsiella* spp. strains used in this study. The STs of KP, as well as KV and KQ included in all analyses are summarized and the number of assembled genomes used in the large pangenome analysis to generate the large patho-core genome is shown. For STs that were further used for omics analysis, information on the representative isolate and its source and original description [26], [42],[43] is provided.

### 2.2. Cultivation and harvest of bacterial strains

All cultivations of bacterial strains were performed in synthetic human urine medium (ULM [originally designated SHU]), composited according to Abbott et al. [44] with shaking (150–220 rpm) at 37°C. For growth kinetics, 5 mL of ULM were inoculated with a single colony of the respective strain (**Table 1**) and incubated overnight. Then, 10 mL of ULM were inoculated at an optical density of 0.05 at λ = 600 nm (OD_600_) and growth was monitored by measurements every 30 min for 8 h. To mitigate the variability in biological replicates (BR), total protein and RNA were extracted from cultures that were initiated through a systematic serial dilution of 5 mL overnight cultures, originating from a glycerol stock. Subsequently, pre-cultures (20 mL of ULM) were inoculated at an OD_600_ of 0.05, starting from mid-exponential phase overnight cultures. Once the pre-cultures reached the mid-exponential phase again, the main cultures (100 mL of ULM) were inoculated in a similar manner. Cell harvest during the mid-exponential phase involved cooling in liquid nitrogen and centrifugation (3 min; 4°C; 8,000 x *g*). Cell pellets were stored at −80°C until RNA and protein preparation.

### 2.3. Isolation of chromosomal DNA and whole genome sequencing

Extraction of total DNA for all strains except PBIO1953 was performed using the MasterPure DNA Purification Kit for Blood, Version 2 (Lucigen, Middelton, WI, USA), according to the manufacturer’s instructions. Afterwards, DNA was quantified with dsDNA HS Assay Kit using Qubit 4 fluorometer (Thermo Fisher Scientific, Waltham, MA, USA) and then shipped to SeqCenter (Pittsburgh, PA, USA). Library preparation was performed using the Illumina DNA Prep Kit and IDT 10 bp UDI indices (Illumina, San Diego, CA, USA), hybrid whole genome sequencing was carried out on an Illumina NextSeq 2000 (2 × 151 bp reads) and Oxford NanoPore on R10.4.1 flowcells run on a GridION platform. Demultiplexing, quality control, and adapter trimming were performed using bcl-convert (v. 3.9.3) [45].

### 2.4. Assembly and annotation

Raw sequencing Illumina reads were adapter-trimmed, and quality-trimmed using Trim galore (https://github.com/FelixKrueger/TrimGalore). Long-reads were assembled with Trycycler (v. 0.5.3) [46] using flye (v. 2.9.2) [47], miniasm and minipolish (v. 0.1.3) [48] and raven (v. 1.8.1) [49]. Resulting genomes were polished with Polypolish (v. 0.5.0) [50] and POLCA (from MaSuRCA v. 4.1.0). In addition, short-reads and long-reads were assembled together with Unicycler (v. 0.5.0). Both assemblies were investigated and merged together, thus we obtained a closed reference with all plasmids. For PBIO1953 the previously assembled closed genome was used. All genomes were annotated with Bakta (v. 1.7.0, database version 5). All genomes were further analysed with kleborate (v. 2.3.2) [26] to verify the ST and determine resistance and virulence features.

### 2.5. Dataset and pangenome analysis

To establish the foundation for subsequent analyses, we curated two distinct genomic datasets. The first dataset included the genomes of two different KV strains (PBIO3543 and PBIO3544) along with eight KP strains (PBIO3546, PBIO3547, PBIO3553, PBIO3554, PBIO3556, PBIO3557, PBIO3559, and PBIO1953). Utilizing the obtained closed genomes of these strains, we constructed a small pangenome. Using the pangenome analysis pipeline (Proteinortho v. 6.2.3) [51], we performed orthologous protein sequence clustering with coverage of ≥50% and e-value 1e^-5^ (other settings: default). The resulting small pangenome was further subdivided into a core genome, containing genes shared by all strains included in the study, and an accessory genome, consisting of genes unique to specific subsets of strains included in the study [52], [53]. The accessory genome was subdivided into the shell genome (genes present in a minimum of two and a maximum of nine strains), the cloud genome (genes unique to a single genome within the dataset), and the patho-core genome (comprising genes shared by all KP strains but absent in KV strains). The visualization of the distribution of genes within the small pangenome was performed using the ggplot2 package and RStudio (v. 4.3.1). For a more detailed understanding of the dataset, we performed a phylogenetic analysis based on whole genome data using mashtree (v. 1.2.0) [54] and visualized with iTOL (v. 6.8.1) and included metadata like ST, resistance score, source of isolation, carriage of ESBL- and carbapenemase-genes, and selected marker genes. Notably, the small pangenome dataset served as a reference for subsequent transcriptomic and proteomic analyses. The second dataset, the large pangenome, included 5,758 publicly available KP genomes from the National Center for Biotechnology Information (NCBI) database [55] (25 Aug 2022), spanning eight globally successful STs, including ST15, ST147, ST258, ST307, along with 982 genomes of KV and KQ, representatives of rather commensal strains. Annotation of genome assemblies was performed using Prokka (v. 1.14.6) [56]. Constructing this large pangenome using cd-hit (v. 4.8.1) [57], we selected cut-offs of 95% identity and coverage to identify potential structural variants. Analogous to the small pangenome, the large pangenome was segmented into a core (present in ≥95% genomes), a shell (present in ≥15%, <95% genomes), a cloud (present in <15% genomes), and a patho-core genome (present in ≥95% KP, <10% KV and KQ genomes). Note that distinct cut-offs were chosen for the large and small pangenomes in orthologous protein sequence clustering based on identity and coverage. This decision was guided by varying sample sizes, the consideration of syntenies during the construction of the small pangenome, and the aim to detect structural variants within the extensive large pangenome.

### 2.6. Isolation of total RNA and sequencing

Total bacterial RNA was isolated by mechanical disruption and acid phenol-chloroform extraction as previously described with minor modifications [58]. The bacterial pellet (equivalent to 8 OD units) was resuspended in 200 µL of ice-cold killing buffer (composition, 20 mM Tris-HCl, pH 7.5; 5 mM MgCl_2_; 20 mM NaN_3_). Subsequently, mechanical disruption was carried out using a Mikro-Dismembrator S (Sartorius, Goettingen, Germany) for 2 min at 2600 rpm. The resulting cell powder was resuspended in 2 mL of lysis buffer (composition, 4 M guanidine-thiocyanate; 25 mM sodium acetate, pH 5.2; 0.5 % (wt/vol) sodium *N*-lauroylsarcosine) prewarmed at 50°C and then stored at −80°C until RNA preparation. The RNA extraction consisted of two washes with 1 volume of acid phenol solution (ROTI^®^Aqua-P/C/I, Carl Roth, Karlsruhe, Germany) and one round with 1 volume 24:1 chloroform-isoamyl alcohol (chloroform ROTIPURAN^®^ ≥99 %/IAA pH 8.0, Carl Roth). The upper phase was transferred into a fresh tube to avoid carry-over of DNA-containing interphase. Following the addition of 1/10 volume of 3 M sodium acetate, RNA was precipitated with 1 mL of isopropanol overnight at −20°C. Upon centrifugation (15 min, 15000 rpm, 4°C) the resulting RNA pellet was washed twice with 70 % (vol/vol) ethanol before the dried pellet was solubilized in 100 µL of RNase-free water. Samples were treated with rDNase (Macherey-Nagel, Düren, Germany) according to the manufacturer’s protocol to minimize DNA contamination. Purification was carried out using the RNA Clean-Up Kit (Macherey-Nagel). Finally, the RNA concentration was determined using a NanoDrop spectrometer (NanoDrop 8000, PeqLab, Erlangen, Germany).

RNA samples were shipped frozen to Novogene (Cambridge, United Kingdom) and, following rRNA depletion and mRNA library preparation, sequenced using 1 × 75 bp reads (Illumina NextSeq 550, nonstranded).

### 2.7. Differential gene expression

Trim Galore (v. 0.6.8) (https://github.com/FelixKrueger/TrimGalore) was used for the adapter and quality trimming of the raw sequencing reads. The trimmed reads were then mapped with Bowtie 2 (v. 2.5.1/mode: – very-sensitive-local) [59], using the assemblies of the previously obtained genomes as references for the individual strains (**Table 1**). The gene counts were calculated using featureCounts (v. 2.0.1/stranded-mode) [60] based on the annotation for the respective strain. Next, the gene names in the count tables were replaced with the cluster IDs of the small pangenome and the different count tables were merged together. The modified count table was imported into RStudio (v. 4.3.1), and differentially expressed genes were identified with DESeq2 (v. 1.40.0) in default mode. The obtained data were further submitted to patho-core analysis.

### 2.8. Isolation of intracellular proteins

Prior to isolation of intracellular proteins, samples were washed twice with ice-cold phosphate-buffered saline (PBS; Thermo Fisher Scientific), and the pellet was suspended in 100 µL of a solution comprising of 20 mM HEPES (pH 8.0) and 2% (wt/vol) sodium dodecyl sulfate (SDS), followed by denaturation at 95°C for 1 min with vigorous shaking. Subsequently, the cells were disrupted using a Mikro-Dismembrator S (Sartorius, Göttingen, Germany) for 3 min at 2600 rpm. The obtained cell powder was then resuspended in 150 µL of preheated (95°C) 20 mM HEPES (pH 8.0) and transferred into a 1.5 mL low binding pre-lubricated tube (Sorenson^TM^ BioScience Inc., Salt Lake City, UT). After cooling to room temperature (RT), 4 mM MgCl_2_ and 0.005 U/µL benzonase (Pierce Universal Nuclease, Thermo Fisher Scientific) were added to the lysates. Ultrasonication was performed for 5 min and the resulting cell debris was pelleted by centrifugation (30 min; RT; 17,000 x *g*). The resulting supernatants containing soluble protein lysates were transferred into a fresh 1.5 mL low binding pre-lubricated tube (Sorenson^TM^ BioScience Inc.) to ensure optimal conditions for further downstream analysis.

### 2.9. Bicinchoninic acid assay

Protein concentration was quantified as previously described [61], [62] using the MicroBCA™Protein Assay Kit (Thermo Fisher Scientific) according to the manufacturer’s instructions. Briefly, samples were diluted at a 1:50 ratio. Concurrently, standards containing a known concentration of bovine serum albumin (BSA) were prepared to match the concentrations of SDS and HEPES in both the samples and standards. All samples and standards were each mixed with equal volumes of the working reaction in duplicates and incubated at 68°C for 1 h. The standards were measured first, followed by sample measurements, all performed in duplicates using the OT-2 robot (Opentrons, Long Island City, NY) and BioTek Synergy (Agilent Technologies, Waldbronn, Germany). The acquired data were evaluated using the Shiny application based on the determined standard curves for accurate quantification [62].

### 2.10. Single pot solid-phase enhanced sample preparation (SP3)

The SP3 protocol was performed as described by Blankenburg *et al.* with slight modifications [61]. For the trypsin digestion, 5 µg of total protein in 10 µL of 20 mM HEPES (pH 8.0) were incubated (18 min shaking at 1400 rpm) with 10 µL of equal volumes of hydrophilic (Speedbead magnetic carboxylated modified particles, GE Healthcare, United Kingdom) and hydrophobic (Sera-Mag Speedbead carboxylated-modified particles, Thermo Fisher Scientific) magnetic beads. The supernatant was discarded by sedimentation of beads on a magnetic rack, followed by two washing steps with 70% (vol/vol) ethanol and one washing step with 100% (vol/vol) acetonitrile (ACN) before air-drying. Trypsin digestion was performed in freshly prepared 20 mM ammonium bicarbonate buffer at trypsin to protein ratio of 1:25 for 18 h at 37°C. The digestion process was stopped by the addition of ACN to a final concentration of 95% (vol/vol) (18 min shaking at 1400 rpm). The beads were further washed and desalted with 100% (vol/vol) ACN and air-dried before elution in 10 µL of 2% (vol/vol) DMSO by sonication for 5 min. The beads were then pelleted by pulse centrifugation, and the resulting supernatant was transferred to a fresh microcentrifuge tube. The eluate was subsequently diluted with an equal volume of buffer A containing 2% (vol/vol) ACN and 0.2% (vol/vol) acetic acid and stored at −20°C until mass spectrometry.

### 2.11. Acquisition and analysis of mass spectrometry data

Peptides were separated using an UltiMate 3000 nanoLC device (Thermo Fisher Scientific) with a pre-column (Acclaim PepMap; Thermo Fisher Scientific) and an analytical column (Accucore; Thermo Fisher Scientific) by applying a binary gradient with buffer A [0.1% (vol/vol) acetic acid in HPLC-grade water] and buffer B (0.1% (vol/vol) acetic acid in ACN) at a flow rate of 300 nL/min. After ionization, peptides were analyzed with a Q Exactive^TM^HF mass spectrometer (Thermo Fisher Scientific) in data independent acquisition (DIA) mode. Specification of the used gradient and detailed settings on nanoLC-MS/MS data acquisition are provided (**Additional file 1: Table S1**). Raw data were mapped against in-house established pangenome database including whole genome sequences of all analyzed strains using Spectronaut (v. 18.1) (Biognosys, Schlieren, Switzerland). All search parameters are provided (**Additional file 1: Table S2**). In brief, trypsin/P was set as the digesting enzyme with 0 missed cleavage allowed. Oxidation at methionine was set as variable modification. Peptides with min. seven amino acids displaying a q-value cut-off for detection of <0.001 were selected for further analyses. Only proteins detected with at least two peptides were considered for statistical analysis, which was performed using RStudio (v. 4.1.3 (2022-03-10)) with the tidyverse package (v. 1.3.2). Normalization factors were obtained from Spectronaut analysis. Missing values (intensity = 0) were replaced with the half-minimal intensity value from the whole dataset. Detected methionine oxidized peptides were excluded from further quantitative analysis. Ion intensities per sample and peptide were summed to calculate peptide intensities. Protein intensities were calculated in the Spectronaut software as intensity-based absolute quantification (iBAQ) values. With the iBAQ algorithm, the summed intensities of the peptides of one protein are divided by the number of theoretically observable peptides (summarized in [63]). iBAQ intensities were separated into quantiles by regarding intensities in the lower two quantiles as close to detection limit. Statistical comparison was accomplished on peptide level with the PECA package (v. 1.30.0) by applying a ROTS test (ROPECA approach - reproducibility-optimized peptide change averaging) [64]. Proteome data have been stored at the ProteomeXchange Consortium via the PRIDE partner repository [65] with the dataset identifier PXD047744. The reviewer account details are: Username, Password: uI1YrOBv.

### 2.12. Patho-core analysis

The patho-core analysis, conducted at genomic, transcriptomic, and proteomic levels, aimed to identify targets that are uniquely present or induced in ULM in all scrutinized pathogenic KP strains compared to their commensal counterparts, KV or KQ (**Table 1**. At the genomic level, we constructed the patho-core genome by filtering for genes present in >95% of each KP ST genome and in <10% of the KV genomes (**Additional file 2: Table S3**). The evaluation of transcriptomic and proteomic data based on the small pangenome followed a systematic approach. Transcriptomic and proteomic profiles of each KP strain were compared individually with each KV strain. In addition, comparisons were made among the KV strains themselves. A gene was considered differentially expressed if the absolute value of the log2 fold change (L2FC) was equal to or greater than 1.5, and the adjusted p-value was less than or equal to 0.05 (|L2FC|≥1.5; p-value adjusted≤0.05). Subsequently, genes were filtered to include only those differentially expressed in all KP strains when compared to both KV strains, excluding those differentially expressed among the KV strains. The cumulative set of genes meeting these criteria was denoted as the patho-core transcriptome or proteome (**Additional file 2: Table S3**). This encompasses genes that may also be present in the genome of KV but are not expressed in ULM, as well as genes absent in KV genomes, yet appearing differentially expressed under infection-mimicking conditions. The different transcriptomic and proteomic profiles were visualized using the pheatmap package in RStudio (v. 4.3.1).

### 2.13. In-depth analysis

For in-depth analysis, genes within the patho-core underwent examination regarding their physiological functions concerning bacterial fitness and virulence. This assessment incorporated Bakta and eggNOG (v. 2.1.9) annotations [66], [67]. The Kyoto Encyclopedia of Genes and Genomes (KEGG) [68] was used for dissecting the involvement of specific genes within their respective functional pathways. Information about the cluster of orthologous groups (COG) was plotted for genes up- or downregulated on transcriptomic or proteomic levels. However, COG annotation could not be determined for all genes. To compensate for this limitation, our functional analysis was expanded through the creation of specialized functional categories. These categories were crafted based on the distinctive characteristics of all regulated genes within the patho-core transcriptome and proteome. The determination of these characteristics drew from information sourced from the aforementioned databases, ensuring that each gene was systematically classified into only one of these categories. This approach not only provided a holistic understanding of the functional landscape but also facilitated the identification of unique features within the patho-core. Additionally, we went beyond typical categorization, closely studying how selected genes relate to bacterial virulence, fitness, and alternative targeting. This exploration aimed to identify specific targets, and to uncover potential areas for further investigation in the context of anti-virulence strategies.

### 2.14. Taxonomy

Representative protein sequences were subjected to a comprehensive BLASTP search against the complete NCBI non-redundant (nr) database to assess the taxonomic distribution of selected markers [55], [69]. This step aimed to elucidate the presence and prevalence of the markers across diverse taxa. Taxonomic identification was pursued at the species level whenever possible; however, in cases where species-level information was unavailable, taxonomic assignments were made at the genus level. In line with the methodology employed for large pangenome clustering, a stringent approach was adopted to exclude structural variants. A cut-off of 95% coverage and identity was applied to the BLASTP search.

### 2.15. Statistical analysis

All phenotypic experiments were performed with three or more independent biological replicates and respective statistical analyses were performed using GraphPad Prism (v. 9.3.0) for Windows (GraphPad Software, San Diego, CA, USA). Data were expressed as mean and standard deviation. Assessment of statistical significance was performed via one-way analysis of variation (ANOVA) with uncorrected Fisher’s LSD. Values lower than 0.05 were used to show significance (*0.033; **0.002; ***<0.001). For transcriptomic and proteomic analyses, a |L2FC| >1.5 with an adjusted p-value equal or smaller than 0.05 was considered as significant.

## 3. Results

### 3.1. Phylogenetic and pangenome analysis revealed differences among KP and KV strains, but also similarities

To identify unique markers, which include genes but also transcripts and proteins, and which might serve as potentially new therapeutical targets in KP, we selected a set of ten *Klebsiella* strains from our strain collection for multi-omics investigation. It consisted of two strains each of four different, successful, international high-risk KP STs, and two different STs of KV which we, based on literature, consider less pathogenic [13], [29].

First, we investigated the distribution of genes among all genomes of the selected dataset by performing a pangenome analysis using Proteinortho (**Figure 1A**). This revealed a total number of 8,519 genes within the pangenome, from which 48% were shared by all strains (core genome). The shell genome consists of genes shared by at least two and a maximum of nine strains and accounts for 30% of the gene content. The cloud genome included 1,784 genes, which were uniquely present in only one strain (approximately 21%). Finally, the most interesting result was a small patho-core genome (SPCG) including 107 genes (1.3%) present in all KP but absent in the KV strains (**Additional file 2: Table S3**). According to the KEGG database, genes within this SPCG were associated with diverse functions in metabolic pathways, encompassing carbohydrates, lipids, nucleotides, amino acids, and other secondary metabolites. We found an enrichment of gene functions related to membrane transport mechanisms, such as ABC transporters, the phosphotransferase system (PTS), and the two-component system. Among others, respective genes encoded proteins of the cellobiose PTS, ensuring uptake and utilization of this carbohydrate. Another example was *fabG*, which is involved in fatty acid metabolism. As stated above, genes with functions in amino acid, but also nucleotide metabolism were quite prominent within the SPCG including the *phn* gene cluster encoding ABC transporter components enabling the sensing and utilization of phosphonates. The KP strains in contrast to the KV strains also encoded a *pgt* gene cluster of the two-component system responsible for phosphoglycerate transport. In addition to these metabolism-related genes, we found *ompC* encoding an outer membrane porin and *kacAT* encoding a type II toxin-antitoxin as unique genes of the KP strains within the SPCG (**Figure 1B**).

**Figure 1:**
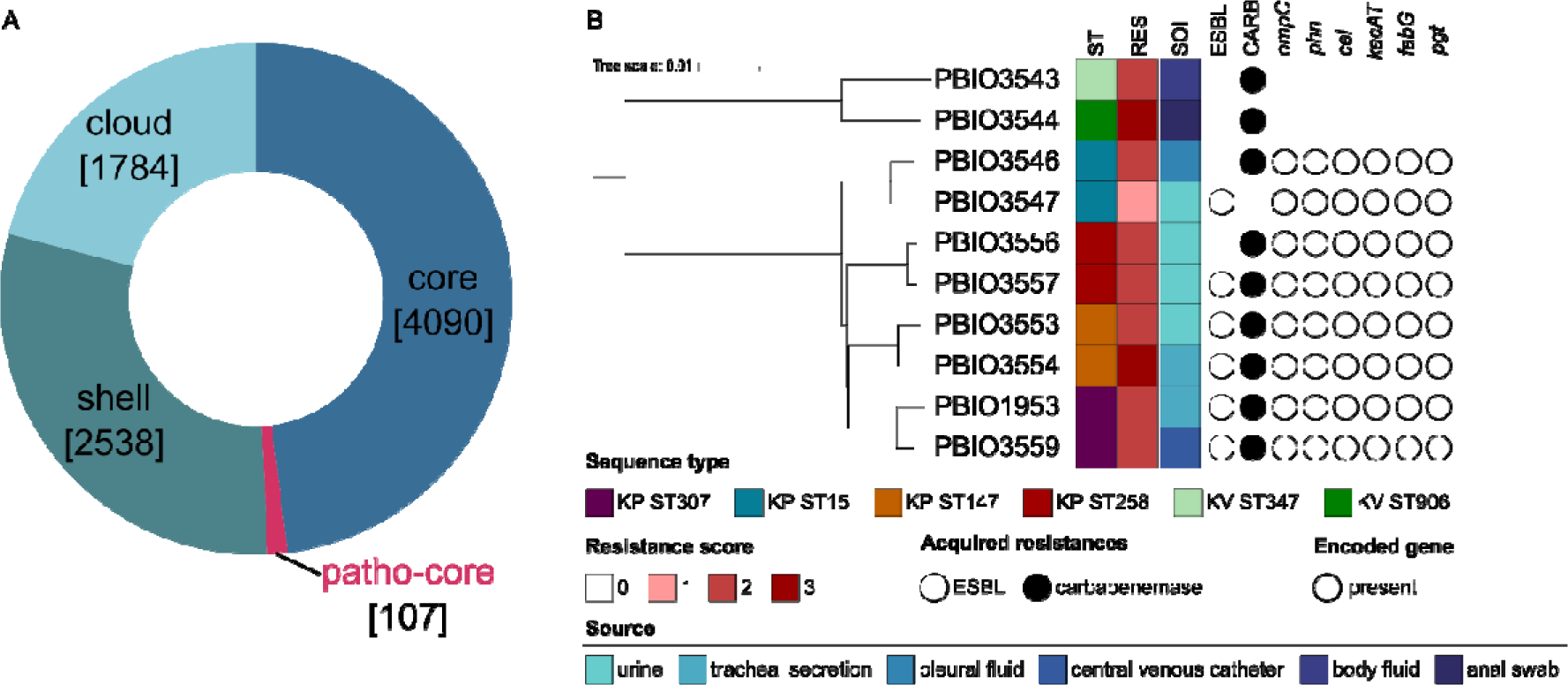
Phylogenetic and pangenome analysis of the selected strains. (**A**) Pangenome analysis grouped genes regarding their occurrence in all strains (core), in minimum two and maximum nine strains (shell), in only one strain (cloud) or in all eight KP but not the KV strains (patho-core). Numbers in square brackets indicate the number of genes within the respective category. (**B**) Based on the whole genomic data, the analyzed dataset including eight different KP and two KV strains was analyzed regarding its phylogenetic relationships considering relevant metadata like the sequence type (ST), resistance score (RES), acquired resistances (ESBL, carbapenemases [CARB]) and the source of isolation (SOI). The phylogenetic tree was constructed using iTOL (v. 6.8.1) and edited with Inkscape.

As expected, the phylogenetic analysis revealed differences between KP and KV and showed a clear separation of the KV from the KP genomes, with the latter closely clustering together, including the respective STs. More importantly, Kleborate revealed a resistance score of >2 for all except one strain (PBIO3547). In addition to *ampC,* each strain carried a minimum of one additional βLJlactamase gene. In five of eight KP strains, genes encoding for both ESBL and carbapenemases were detected. Note that all strains originate from human sources with KV strains obtained from body fluid and an anal swab (PBIO3543 and PBIO3544, respectively). Four KP strains were isolated from urine (PBIO3547, PBIO3556, PBIO3557, and PBIO3553), two from tracheal secretion (PBIO3554, and PBIO1953), and one from pleural fluid (PBIO3559).

In summary, phylogenetics combined with genomic analysis of a selected *Klebsiella* dataset demonstrated differences between the two phylogroups, impacted by the presence of genes supporting metabolic features in KP in contrast to KV.

### 3.2. Patho-core transcriptome and proteome analysis in infection-mimicking conditions demonstrated similar regulatory patterns among KP in contrast to KV strains

Following the genomic analysis, which revealed 107 patho-core genes in KP, we next screened for differentially expressed genes and proteins to i) explore whether the KP patho-core genome would translate to the transcriptome and/or proteome and ii) to identify different core markers and regulatory patterns on transcriptomic and/or proteomic levels. Given that KP is a leading cause of UTI, we generated our samples in ULM, which likely reflects relevant infection-mimicking conditions. Interestingly, KP grew significantly better in ULM than the KV strains (*p* <0.001) (**Figure 2A+B**). However, this significant difference between KV and KP strains was not observed when grown in LB medium (**Additional file 1: Figure S1**). Together with the above-mentioned genomic data (**Figure 1B**), this suggests a better utilization capacity of nutrients by the KP strains. This not only applied to the strains obtained from UTI but all KP representatives.

**Figure 2:**
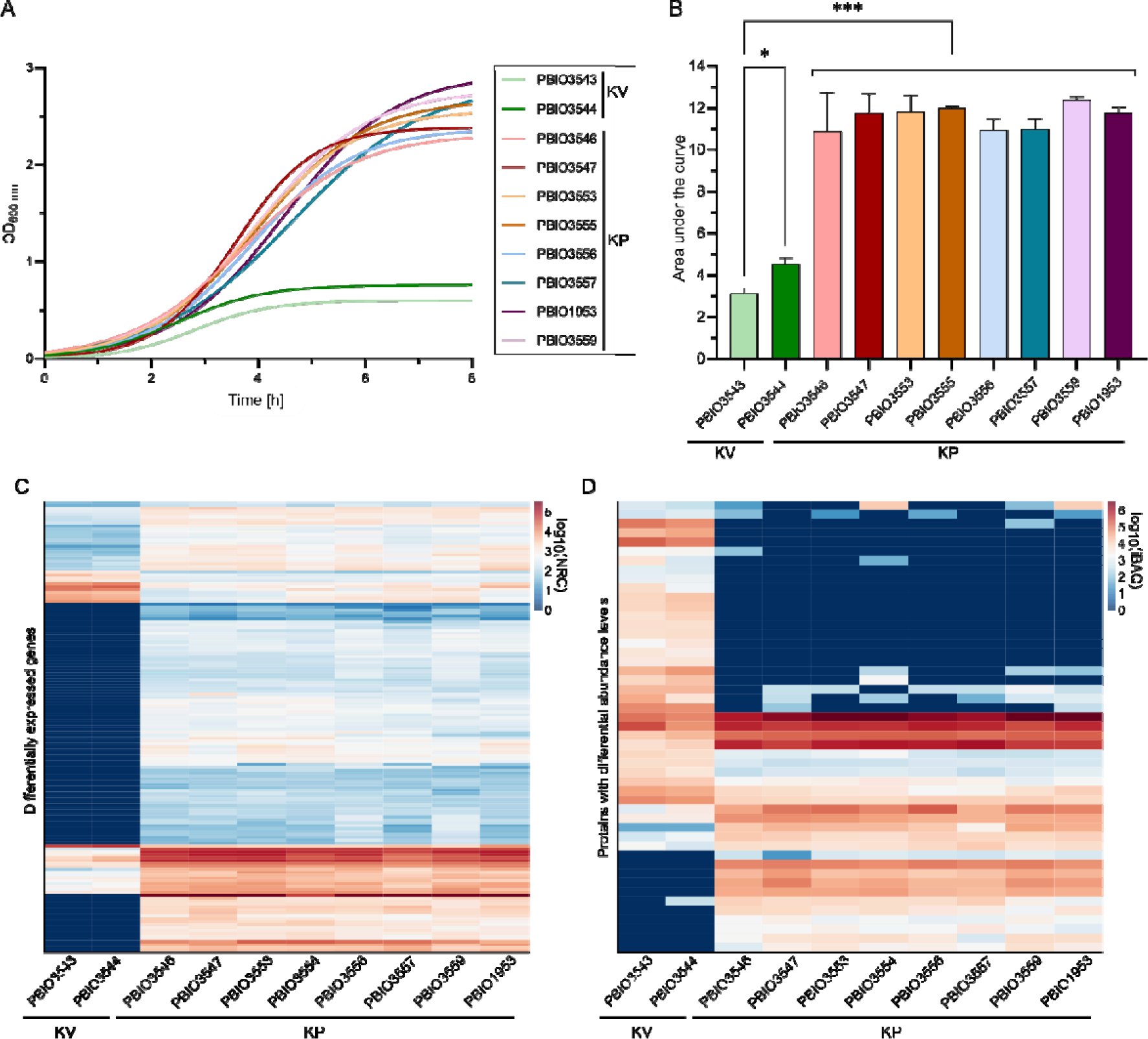
Growth kinetics and expression profiles of KP and KV strains in ULM. Strains were cultivated in ULM in biological triplicates. (**A**) Growth kinetics are displayed using GraphPad Prism and Gompertz-Growth fitting. (**B**) The area under the curve was calculated from three individual cultivations (n=3). The significance of differences was calculated using one-way ANOVA (p-value: * 0.033; *** <0.001). (**C+D**) The normalized read counts obtained from transcriptomic analysis (**C**) and the iBAQ obtained from proteomic analysis (**D**) are plotted for each strain. The regulatory profiles of genes within the patho-core transcriptome or proteome are shown as separate heatmaps. The log10 was applied to all values for better visualization of differences.

Then, transcriptomic and proteomic analyses were performed for all eight KP and two KV strains after cultivation in ULM to explore the underlying processes potentially supporting enhanced growth of KP. We observed differences among all KPs and KVs at both transcriptomic and proteomic levels, but also differences among the KP strains, while strains of the same ST clustered closely together (**Additional file 1: Figure S2**). Therefore, we compared the data of each KP with each KV strain and the two KV strains with each other, while only considering distinct differences (|L2FC|≥1.5; p-value adjusted≤0.05).

By plotting the decadic logarithm (log_10_) of the normalized read counts (NRC) and the log_10_ of iBAQ values regulatory patterns of genes within the patho-core transcriptome and proteome revealed that overall patterns were seemingly different on transcriptomic versus proteomic levels (**Figure 2C+D**). However, they clearly show that all KP strains shared a similar expression profile, which was different from the two KV strains.

Genes that appeared significantly expressed in all comparisons of KP vs. KV strains, but not between the two KV strains, were assigned to the patho-core transcriptome (PCT) or proteome (PCP), respectively. In general, the PCT contained a larger number of genes than the PCP (**Additional file 1: Figure S3**). Moreover, a higher number of genes was upregulated in all individual comparisons of KP vs. KV on transcriptomic levels. Upregulated genes within the PCT included 143, while only a total number of 12 genes were downregulated. Up- and downregulations on proteomic levels were rather equal for the individual comparisons, with a slightly higher number of downregulations (29) than upregulations (20) within the PCP. The total number of regulated genes suggests that the strains responded highly individually to the specific growth conditions of ULM. Nevertheless, a substantial patho-core transcriptome and/or proteome indicates common response mechanisms in KP, which were absent in KV.

Among the 155 genes identified within the PCT, 103 were assigned to the SPCG, while 52 were found in the core genome of the selected *Klebsiella* dataset. Of the 49 genes belonging to the PCP, six were discovered in the SPCG, 24 were assigned to the core, and 19 to the shell genome of the selected dataset. This multi-omics analysis highlights that almost all genes of the SPCG exhibited transcriptional activity, with detectable RNA levels under conditions mimicking urine. However, it also suggests that only a minority undergoes sufficient translation to be detected by proteomics under these experimental conditions. In addition, we identified genes from the core genome that exhibited unique expression patterns in KP strains, thereby being integral components of the PCT and/or PCP. This possibly means that growth advantages of KP strains in ULM are not solely attributed to their particular genetic composition but are also substantiated by shared regulatory and response mechanisms that contribute to overall fitness.

### 3.3. In-depth characterization revealed common functional pathways on transcriptomic and proteomic levels in KP

The observed regulatory profiles of the patho-core on transcriptomic and proteomics levels for KP were further investigated regarding their physiological functions. Since categorization based on the COG database was insufficient (**Additional file 1: Figure S4**), specialized categories were created (**Additional file 3: Table S4**). Genes were first differentiated based on their up- or downregulation compared to the respective gene in KV and then clustered into four different groups, namely porin/transporter, metabolism, regulation/stress response, and others (**Figure 3**). Then, for a more detailed characterization, smaller categories were assigned for all regulated genes (**Additional file 4: Table S5**). Indeed, transcriptomic and proteomic profiles not only varied in their number of up- or down-regulated genes but also in their putative functions. Nevertheless, some similarities were found. For instance, regulation of metabolic pathways was seemingly a key mechanism when growing KP in ULM. On transcriptomic levels, 49% of the up- and 25% of the downregulated genes were assigned to this group, while on proteomic levels, 65% of proteins with higher abundance in KP compared to KV and 41% of proteins with lower abundance in KP compared to KV were involved. Moreover, most of the more specific categories were shared on both transcriptomic and proteomic levels, namely sulfur, nitrogen/ammonia, citrate, phosphate, biotin/fatty acid, carbohydrate, amino acid, ribosomes/tRNA, and peptide metabolism. Downregulations also affected transporters, with 25% on transcriptomic and 21% on proteomic levels. Genes with regulatory functions and involved in stress response had a greater impact on the proteomic (10% up- and 17% downregulated, respectively) than on the transcriptomic level (12.5% up- and 8% downregulated, respectively). Nevertheless, both up- and downregulation of other regulatory genes seemed to be common.

**Figure 3:**
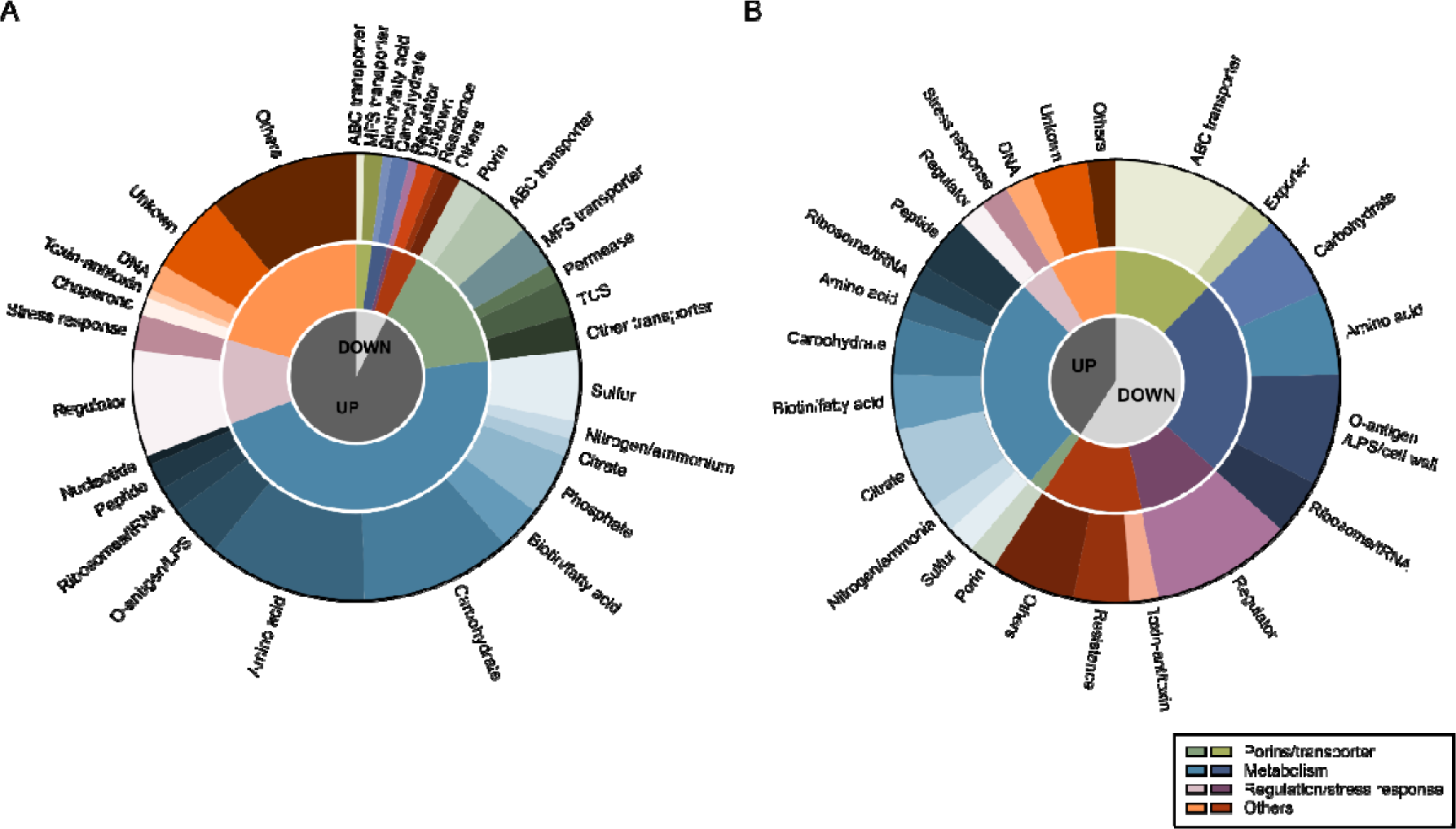
Functional analysis of regulated genes of the PCT and PCP. Functional evaluation of all genes within the PCT (**A**) and PCP (**B**) was performed by creating categories on three different levels of specificity based on all regulations. First, we only considered fold changes (L2FC) >1.5 (upregulated [UP]) or <-1.5 (downregulated [DOWN]). Second, four different groups were assigned for broad categorization: porins/transporter, metabolism, regulation/stress response, and others, as indicated in the legend. Third, more specific categories within the previously differentiated groups were chosen and each of the PCT and PCP was assigned to these categories. Visualization was done using GraphPad Prism and Inkscape.

Downregulation of genes involved in resistance was observed on both levels, while genes for DNA replication and repair were upregulated. Regulations among the levels also varied concerning toxin-antitoxin systems and chaperones. Overall, the in-depth characterization revealed that the regulations on a functional level are rather similar on transcriptomic and proteomic levels and that they mainly concern metabolic pathways.

### 3.4. Comparing regulated genes to a large pan- and patho-core genome of the *Klebsiella* genus identified unique markers in KP

The 193 differentially expressed genes on both transcriptomic and proteomic levels between KP and KV in ULM were then further investigated to explore whether they would play a role as unique markers in a broader context. We therefore leveraged >6000 publicly available *Klebsiella* spp. genome assemblies comprising KV and KQ as less successful representatives and eight different, successful international high-risk KP STs (i.e. ST11, ST15, ST23, ST101, ST147, ST258, ST307, and ST512) (**Additional file 1: Figure S5**) to construct a large pangenome (not to be confused with the above mentioned “small” pangenome) (**Additional file 2: Table S3**).

All proteins were clustered based on greater than and equal to 95% identity and coverage, to also decipher structural variants. The resulting pangenome consisted of 192,172 clusters within the cloud genome and 2,785 within the shell genome. Within the core genome, we detected 3,083 clusters, while 273 clusters were classified as patho-core genes. We based this large patho-core genome (LPCG) on the cut-offs that clusters were present in a maximum of 10% of each of the KV and KQ genomes and a minimum of 95% of each KP ST. We identified genes that were (i) detected within the LPCG and differentially expressed in the transcriptome and/or proteome, (ii) differentially expressed (either on transcriptomic and/or proteomic levels) but did not match the criteria for being included in the LPCG, and (iii) detected within the LPCG, but were not differentially expressed at all (**Additional file 1: Figure S6**). Overall 31 genes were part of the LPCG and either regulated on transcriptomic and/or proteomic levels (i), while most genes were not included in the LPCG but detected either as part of the patho-core transcriptome and/or proteome (ii). This suggests that the functionally investigated KP strains share common gene contents and regulatory mechanisms leading to the expression of relevant genes in specialized settings such as ULM or the urinary tract. Regardless of their distribution within the pangenome and whether they belonged to the large patho-core genome, we analyzed all 193 differentially expressed genes on transcriptomic and/or proteomic levels regarding their biological relevance using different databases such as UniProt, KEGG, STRING, and considered previous publications. This demonstrated that, although genes were present in all *Klebsiella* spp. genomes and thus part of the large pangenome, unique induction occurred in KP in urine-like conditions. As mentioned above, most of the regulations addressed metabolic mechanisms (**Figure 3**). In agreement with that, during the in-depth analysis, genes with potential metabolic functions were predominantly characterized with a higher biological relevance (**Table 2**), indicating the overall metabolic benefit of KP compared to KV under urine-like conditions. As the most interesting example for the first category (i), we identified genes for a cellobiose PTS (*celABF*) that were significantly expressed on transcriptomic and/or proteomic levels in urine-like conditions and absent in KV/KQ while present in the majority (≥95%) of all KP genomes (**Figure 4**). In addition, these genes were also detected within the SPCG. It is well-known that bacteria encode several variants of cellobiose-specific uptake systems [70]. Hence, we investigated the presence of gene variants of *celB* in a representative strain (PBIO1953). This showed that genes with similarity to *celB* occurred to a maximum of 30% identity. Another interesting candidate of this category was *fabG*, which has a central function in fatty acid biosynthesis. In the second category (ii), genes involved in citrate uptake were identified. While they occurred equally in all *Klebsiella* spp. genomes, they were only induced in KP (**Figure 4**). Citrate is highly abundant in urine and strains encoding for specific citrate uptake genes are able to use it as a sole carbon source under anaerobic conditions [71]. Another group of genes found in the second category (ii) and which was present in >99.8% of the investigated genomes, encodes for a cytochrome ubiquinol oxidase (*cydAB*), which enables coupling the reduction of molecular oxygen to the generation of a proton motif force and thus energy production. In addition, on transcriptomic levels, we found two gene clusters upregulated in the KP strains, which are responsible for the acquisition of different phosphorus sources. The first cluster included seven genes of a phosphonate ABC-transporter (*phnRXVTWSU*). While these genes were missing in the majority of the KV genomes, most KQ and KP genomes carried them. Similar applied to the second gene cluster encoding for genes for a two-component system involved in phosphoglycerate transport (*pgtABCP*). This gene cluster did not occur in KV but in 10–50% of the KQ genomes and in more than 92% of the KP genomes.

In summary, we identified several unique markers in KP belonging to the different categories.

**Table 2:**
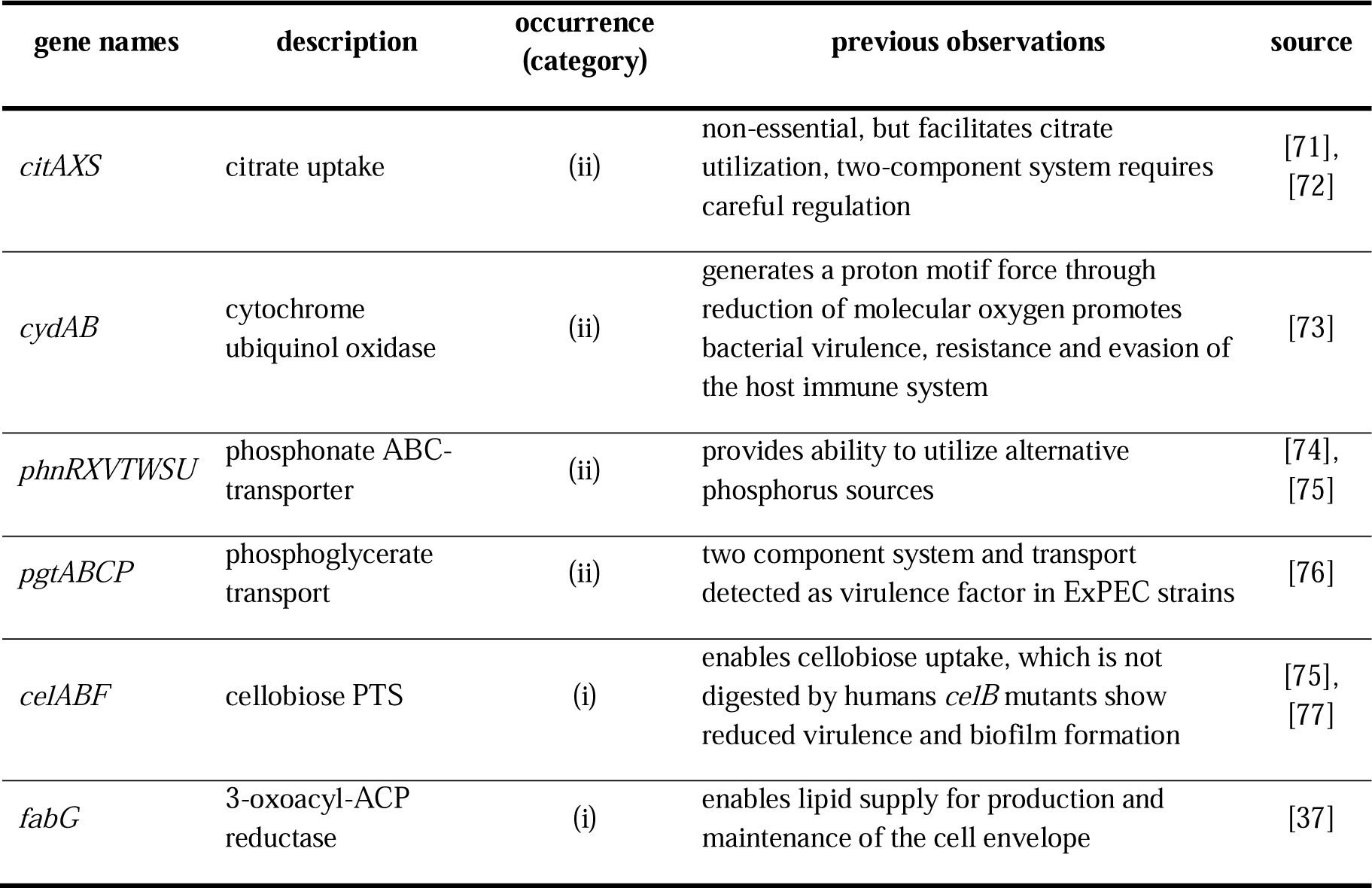
Selected differentially expressed genes with particular biological relevance. The gene names, a description, as well as summary of previous observations and the respective sources are shown for selected groups of genes which resulted from the in-depth multi-omics analysis.

**Figure 4:**
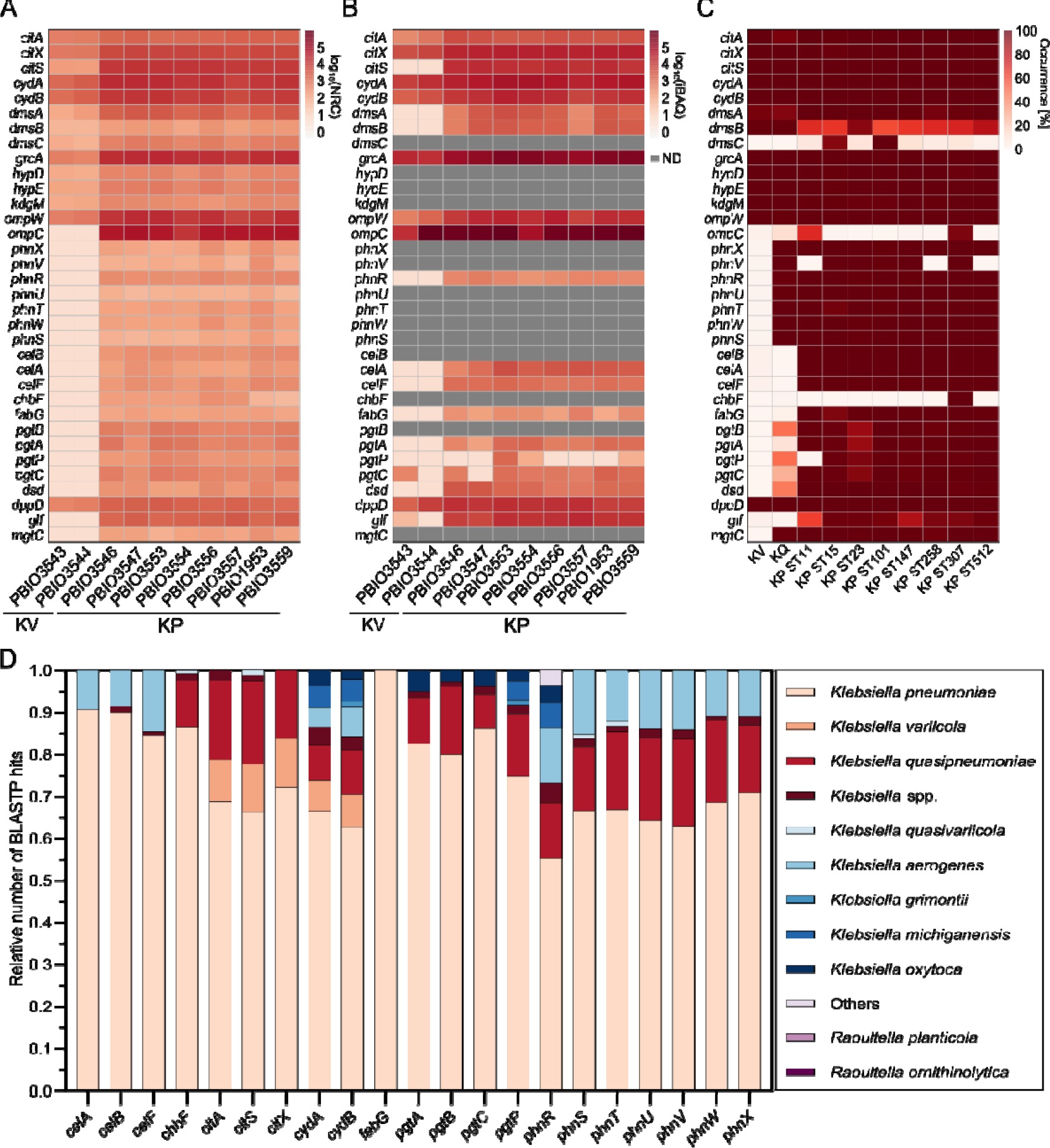
Detection of corresponding gene expression on multi-omics levels and taxonomic distribution of the selected markers. Genes identified as differentially expressed on transcriptomic and/or proteomic levels were selected regarding their biological relevance based on literature research. Heatmaps for all profiles, either expression on transcriptomic or proteomic level or occurrence on genomic level are shown and respective genes, strains or STs are indicated. ND means the protein was not detectable. (**A**): Transcriptomic profile of selected genes was generated by mapping the log10 of the NRC for each analyzed strain. (**B**): Profile of the expression detected on proteomic level of selected genes was generated by mapping the log10 of the iBAQ values for each analyzed strain. (**C**): Selected genes were mapped against the LPCG (BLASTP). (**D**): The protein sequences representing the indicated markers were subjected to BLASTP analysis against the NCBI nr database, employing a stringent cut-off of 95% coverage and identity. Taxonomic identification was pursued at the species level whenever feasible; otherwise, results were reported at the genus level. The figure illustrates the number of hits observed for each selected marker across various species. Species with fewer than five hits for a particular marker were consolidated under the category “Others.”

### 3.5. Analysis of the taxonomic distribution of selected markers revealed their predominance in the *Klebsiella* genus

The distribution of the respective protein sequences corresponding to each selected marker (**Table 2**) was examined across various species and genera using the NCBI nr database and BLASTP. To ensure a robust dataset, a filtering criterion was applied, setting a cut-off of 95% identity and coverage. This approach yielded approximately 160 to 1,200 hits for each representative sequence (**Additional file 5: Table S6**). Plotting markers with more than five hits in one species revealed that the majority of hits was prominently associated with KP (**Figure 4D**). It is crucial to acknowledge a potential bias stemming from the disparate number of genomes included in the NCBI database for KP. Notably, our analysis underscored that, beyond KP, the *Klebsiella* genus as a whole exhibited substantial prevalence. Other species within this genus included KV, KQ, *K. aerogenes, K. africana, K. grimontii, K. michiganesis, K. oxytoca, K. pasteurii, “K. quasivariicola”, K. spallanzanii,* and a minority of other *Klebsiella* species. Moreover, in 17 of 21 of the selected markers, we identified species belonging to genera other than *Klebsiella*. These non-*Klebsiella* genera exhibited a minority presence, with less than six hits (<2%) for each marker including *Citrobacter portucalensis, Clostridium perfringens, Enterobacter* spp.*, Escherichia coli, Kluyvera ascorbata, Kluyvera cryocrescens, Kluyvera intermedia, Kluyvera* genomosp. 1*, Kluyvera georgiana, Pseudocitrobacter corydidari, Raoultella ornithinolytica, Raoultella planticola, Raoultella* spp.*, Salmonella enterica, Samsonia erythrinae,* and *Veillonella dispar*.

## 4. Discussion

Here, we applied a multi-omics approach to identify unique markers in *K. pneumoniae,* an often MDR pathogen predominantly occurring worldwide [1]. By generating transcriptomic and proteomic data for a set of clinical *Klebsiella* isolates and including more than 6,000 publicly available whole genomes, we identified several potentially novel and KP-specific targets [78]. Multi-level omics studies reveal adaptive and regulatory mechanisms that likely contribute to the success of certain pathogens in different ecologies and over other, rather opportunistic, bacteria [78], [79]. However, measuring general and clinical “success” is very complex and depends not only on the individual infection context but on the broad distribution and bacterial characteristics such as AMR and virulence [29], [80]. For this study, we compared KP strains that belong to acknowledged successful, international high-risk clonal lineages with putatively less successful KV (and KQ) representatives isolated from different clinical sources [12], [13]. We hypothesized that KP harbors unique markers absent or not induced in KV/KQ, which could be prospectively leveraged as novel antimicrobial targets. Unique markers might occur because successful KP lineages have undergone adaptive evolutionary processes that resulted in core genes or gene functions that are different from less successful bacteria [13], [29].

Indeed, for our strains, growth kinetics in urine-like conditions showed that KP likely utilizes a broader spectrum of nutrients, leading to improved growth and overall adaptation to this particular surrounding, when compared to KV. This was supported by the unique presence of various gene clusters (e.g., *phn, pgt,* and *cel*) within the SPCG associated with improved metabolism. It is of course well-known that adequate response and adaptation to specialized host environments such as the urinary tract, is a prerequisite for the establishment of a following infection [6], [81]. Subsequent proteome and transcriptome analyses supported our initial findings. Not only did we detect similar regulatory patterns for all KP in comparison to the KV strains but our results underline the assumption that induced metabolic features enable improved responding to specialized environments. These differentially expressed metabolic pathways included citrate acquisition, the cytochrome ubiquinol oxidase, phosphorus source- and cellobiose utilization, as well as fatty acid biosynthesis, and are seemingly important for general survival and growth [73], [77], [82], [83].

Upscaling of our finding to a larger *Klebsiella* data set (LPCG) revealed different unique markers in KP vs. other *Klebsiella*. For example, we detected genes encoding for the cellobiose PTS on all omics levels. Association of PTS and virulence has been previously shown for the connection of the fructose PTS and fimbriae expression in *E. coli* [84]. Also, the induction of the bacterial _L_-fructose metabolism seemingly promotes gastrointestinal colonization by KP, which is a prerequisite for infection [85], [86]. In addition to host colonization and adaptation, bacteria compete with other microorganisms for nutrients, which is likely due to contrary preferential usage of nutrients [87]. This broad metabolic flexibility, also described for KP, is necessary for the ability to cause multiple site infection in hosts [6]. Metabolic adaptation has been described for extraintestinal *E. coli,* which need to switch their metabolism when changing from a commensal lifestyle within the intestine to a pathogenic lifestyle within the urinary tract [81]. Interestingly, it has been previously shown that metabolic adaptation through, e.g., two-component systems can trigger the expression of virulence-associated genes [81]. In our case, this could be true for the induction of the cellobiose PTS including *celB*, which is involved in bacterial biofilm formation and *in vivo* virulence in a mouse model of intragastric infection [77].

Markers that were present in KQ and KP genomes but absent KV included genes for the phosphor metabolism (*phn and pgt*). The broad occurrence of *phn* genes in clinical KP isolates, important for 2-aminoethylphosphonate transport and degradation, has already been reported [74], [75]. Another example of the association between metabolism and virulence in KP under urine-like conditions is the *pgt-*operon, which is not only responsible for phosphoglycerate uptake via a two-component system but encoded on a pathogenicity island in uropathogenic *E. coli* together with the virulence-associated capsule (*kps*) locus (K15) [7], [88].

Interestingly, we detected KP-specific induction of the *cydAB* genes in ULM, although the genes were present in all investigated bacterial species and in more than 99.8% of the genomes. Besides the above-mentioned generation of proton motif forces, several other implications have been previously reported. This includes not only the involvement in stress response, virulence, and bacterial evasion of the host immune system but also their role in resistance to several classes of antimicrobials [73]. A gene cluster also found to be equally distributed among KV, KQ, and KP genomes, but only induced in KP in ULM, contributes to citrate uptake. Note that citrate is abundant in urine, making this inducible response particularly relevant in UTI [89]. Interestingly, it has been previously shown that genes necessary for citrate uptake not only provide the ability for citrate utilization but also environmental sensing, and the tight regulation of this process by the two-component system CitAB [72].

Cells modulate gene expression in response to environmental changes, particularly in the context of nutrient depletion and availability [90]. Hence, in the context of an *in vitro* condition in a nutrient-defined artificial urine medium, we expected the expression of genes involved in specialized metabolic pathways including the utilization of the nutrients present. However, it also has been frequently reported in *in vivo* settings that metabolic pathways play a major role in KP infection [37], [91]. For example, by using a transposon library, Bachman and colleagues elucidated key processes crucial for KP fitness in a lung infection model, pinpointing genes linked to bacterial metabolism, such as branched-chain amino acid metabolism [91]. The importance of metabolic flexibility has been demonstrated for mucosal surface colonization even prior to the establishment and manifestation of infection [81], [83], [90]. Our results suggest increased metabolic flexibility of KP vs. other *Klebsiella* in urine-like conditions, which possibly contributes the success of KP as an opportunistic pathogen. This is underlined by the detection of genes, e.g., the *cit* gene cluster, among all *Klebsiella* genomes, but the KP-specific induction in ULM. Moreover, these results suggest that KP responds to the ULM environment by regulating a diverse set of genes, some of which have the potential to be targeted for therapeutic interventions. Such interventions might be implemented in the context of anti-virulence strategies for instance by the individual dysregulation of respective metabolic pathways [92]. An example for the latter is the folate metabolism addressed by trimethoprim, which results in bacterial stress due to the lack of key metabolites important for DNA, RNA, and protein synthesis [93]. As mentioned above, we revealed unique induction of the two-component system CitAB in KP vs. KV in ULM. The tight regulation provided by CitAB, which senses and regulates citrate utilization, could be potentially disrupted to specifically decrease KP fitness and thus prevent infection manifestation [12], [72]. Another interesting candidate for anti-virulence approaches is FabG, as other proteins (e.g., FabI, FabB, FabH) of this pathway have already been described regarding potential KP drug targets [37].

In contrast to conventional antibiotics, anti-virulence targets offer the advantage of minimizing potential side effects on especially crucial commensal bacteria and lower resistance development [39].

## 5. Conclusion(s)

In conclusion, our study provides valuable insights into the genomics and adaptive responses of KP, particularly under infection-mimicking conditions. Understanding the expression of key genes and pathways unique to pathogenic KP vs. other *Klebsiella* strains contributes to the development of anti-virulence strategies and potential therapeutic targets in the framework of “precision medicine”. Prospective studies will have to explore further, whether the identified markers in this study truly hold up as druggable novel targets.

## 6. Declarations

### 6.1. Ethics approval and consent to participate

Where applicable, ethical approval was given by the ethics committee of the University Medicine Greifswald (BB 133/20). Informed patient consent was either waived as samples were taken under a hospital surveillance framework for routine sampling, or as samples were sent to the German national reference laboratory. The research conformed to the principles of the Helsinki Declaration.

### 6.2. Consent for publication

Not applicable

### 6.3. Availability of data and materials

The experimental and computational data that support the findings of this research are available in this article and its supplementary information files. Genomic and transcriptomic data for this study have been deposited in the European Nucleotide Archive (ENA) at EMBL-EBI under accession number PRJEB71341 (https://www.ebi.ac.uk/ena/browser/view/PRJEB71341).

Proteome data have been stored at the ProteomeXchange Consortium via the PRIDE partner repository [65] with the dataset identifier **PXD047744.** The datasets used and/or analyzed during the current study are available from the corresponding author on reasonable request.

### 6.6. Authors’ contributions

Conceptualization K.Sc.; methodology, K.Su., L.S., and M.S.; software, K.Su., L.S., M.S. and S.E.H.; validation, L.S.; formal analysis, J.M., K.Su., L.S., and M.S.; investigation, K.Su., L.S., M.S., and M.S.G; resources, G.W., J.B, K.B., K.Sc, N.H., S.G. and U.V.; data curation, K.Su, L.S., M.S. and S.E.H..; writing-original draft preparation, L.S.; writing-review and editing, E.E., K.Su, and K.Sc,.; visualization, L.S. and M.S.; supervision, K.Sc.; project administration, K.Sc.; funding acquisition, K.Sc. All authors have read and agreed to the published version of the manuscript.

## 6.7. Acknowledgments

We thank Dr. Stephan Michalik for creating the R scripts for statistical analysis of proteome data and Sara-Lucia Wawrzyniak for her excellent technical support.

## 7. Abbreviations

(ACN): Acetonitrile
(AMR): Antimicrobial resistance
(BR): Biological replicates
(BSA): Bovine serum albumin
(cKp): Classical *K. pneumoniae*
(COG): Cluster of orthologous groups
(log_10_): Decadic logarithm
(ESBL): Extended-spectrum β-lactamases
(hvKp): Hypervirulent *K. pneumoniae*
(iBAQ): Intensity-based absolute quantification
(K).: *Klebsiella*
(KP): *Klebsiella pneumoniae*
(KpSC): *Klebsiella pneumoniae* species complex
(KQ): *Klebsiella quasipneumoniae*
(KV): *Klebsiella variicola*
(KEGG): Kyoto Encyclopedia of Genes and Genomes
(LPGC): Large patho-core genome
(L2FC): log2 fold change
(MDR): Multidrug-resistant
(MLST): Multi-locus sequence typing
(NDM-1): New Delhi metallo-beta-lactamase 1
(NRC): Normalized read counts
(PCP): Oxacillinase-48 (OXA-48) Patho-core proteome
(PCT): Patho-core transcriptome
(PTS): Phosphotransferase system
(SDS): Sodium dodecyl sulfate
(STs): Sequence types
(SP3): Single pot solid-phase enhanced sample preparation
(SPCG): Small patho-core genome
(ULM): Synthetic human urine medium
(UTI): Urinary tract infection

