## Supplementary material for "Multi-omics investigations uncover unique pathogenic markers in clinical *Klebsiella pneumoniae* that could be leveraged as novel antimicrobial targets": Table S1-S2 and supplement figures

Table S1: Instrumental setting for Reversed phase liquid chromatography (RPLC) and mass spectrometry.

| Reversed phase liquid chromatography (RPLC) | |
| --- | --- |
| Instrument | Ultimate 3000 RSLC (Thermo Scientific) |
| Trap column | 75 μm inner diameter, packed with 3 μm C18 particles (Acclaim PepMap100, Thermo Scientific) |
| Analytical column | Accucore 150-C18, (Thermo Fisher Scientific)  25 cm x 75 μm, 2.6 μm C18 particles, 150 Å pore size |
| Buffer system | binary buffer system consisting of 0.1% acetic acid in HPLC-grade water (buffer A) and 100% ACN in 0.1% acetic acid (buffer B) |
| Flow rate | 300 nl/min |
| Gradient | 0 min 2% B 🡪  2 min 5% B 🡪  10 min 5% B 🡪  130 min 25% B 🡪  135 min 40% B 🡪  137 min 90% B 🡪  142 min 90% B 🡪  145 min 2% B 🡪  150 min 2% B |
| Gradient duration | 120 min |
| Column oven temperature | 40°C |
| **Mass spectrometry** | |
| Instrument | Q Exactive HF mass spectrometer |
| Electrospray | Nanospray Flex Ion Source |
| Operation mode | data-independent |
| **Full MS** |  |
| MS scan resolution | 60000 |
| Norm. AGC target | 5e6 |
| maximum ion injection time for the MS scan | 200 ms |
| Scan range | 333 to 1650 m/z |
| RF Lens | 50% |
| Spectra data type | profile |
| **dd-MS2** |  |
| Precursor mass range | 333 to 1650 m/z |
| Resolution | 30,000 |
| Norm. MS/MS AGC target | 3e6 |
| Maximum ion injection time mode | auto |
| Spectra data type | profile |
| Microscans | 1 |
| Isolation window | 56 windows, 13 m/z, 2 m/z overlap |
| Define first mass | 200 |
| Dissociation mode | higher energy collisional dissociation (HCD) |
| Normalized collision energy | 27.5 % |

Table S2: Spectronaut settings for mass spectrometry.

ANALYSIS DATA

Spectronaut 18.1.230626.50606

Analysis Mode: UI

Analysis Type: directDIA

Analysis Date: 27-July-2023 10:50:27 UTC+0

BEGIN-SETTINGS

Settings Used: C_FunGene_directDIA_sparse_no_imputing

├─ DIA Analysis\Calibration

│ ├─ MZ Extraction Strategy:	Maximum Intensity

│ ├─ Allow source specific iRT Calibration:	True

│ ├─ Precision iRT:	True

│ │ ├─ Exclude De-amidated Peptides:	True

│ │ └─ iRT <-> RT Regression Type:	Local (Non-Linear) Regression

│ ├─ MS1 Mass Tolerance Strategy:	System Default

│ └─ MS2 Mass Tolerance Strategy:	System Default

├─ DIA Analysis\Identification

│ ├─ Precursor Qvalue Cutoff:	0.001

│ ├─ Precursor PEP Cutoff:	0.2

│ ├─ Protein Qvalue Cutoff (Experiment):	0.01

│ ├─ Protein Qvalue Cutoff (Run):	0.05

│ ├─ Protein PEP Cutoff:	0.75

│ ├─ Single Hit Definition:	By Stripped Sequence

│ ├─ Exclude Single Hit Proteins:	False

│ ├─ Exclude Duplicate Assays:	True

│ ├─ Exclude Predicted Fragment Scores:	False

│ ├─ Generate Decoys:	True

│ │ ├─ Decoy Generation Method:	Mutated

│ │ │ └─ Preferred Fragment Source:	NN Predicted Fragments

│ │ └─ Decoy Limit Strategy:	Dynamic

│ │	└─ Library Size Fraction:	0.1

│ └─ Pvalue Estimator:	Kernel Density Estimator

├─ DIA Analysis\Pipeline Mode

│ ├─ Generate SNE File:	True

│ │ └─ Store Ion traces in SNE:	False

│ ├─ Post Analysis Reports:

│ │ ├─ CV Density Line Chart:	True

│ │ ├─ CVs Below X Bar Chart:	True

│ │ ├─ Data Completeness Bar Chart:	True

│ │ ├─ Run Identifications Bar Chart:	True

│ │ └─ Scoring Histograms:	True

│ ├─ Report Schema:	C_FunGene_complex (Normal), C_FunGene_complex (Normal)

│ └─ Reporting Unit:	Across Experiment

├─ DIA Analysis\Post Analysis

│ ├─ Differential Abundance Testing:	Paired t-test

│ │ ├─ Group-Wise Testing Correction:	False

│ │ ├─ Log2 Ratio Candidate Filter:	0.58

│ │ └─ Confidence Candidate Filter:	Qvalue

│ │	└─ Confidence:	0.05

│ ├─ Differential Abundance Grouping:	Major Group (Quantification Settings)

│ │ └─ Smallest Quantitative Unit:	Precursor Ion (Quantification Settings)

│ │	└─ Use All MS-Level Quantities:	False

│ ├─ Calculate Explained TIC:	None

│ ├─ Calculate Sample Correlation Matrix:	True

│ └─ Hierarchical Clustering:	True

│	├─ Distance Metric:	Manhattan Distance

│	├─ Linkage Strategy:	Ward's Method

│	├─ Order Runs by Clustering:	True

│	└─ Z-score Transformation:	False

├─ DIA Analysis\Protein Inference

│ └─ Protein Inference Workflow:	Automatic

│	└─ Inference Algorithm:	IDPicker

├─ DIA Analysis\PTM Workflow

│ └─ PTM Localization:	False

├─ DIA Analysis\Quantification

│ ├─ Precursor Filtering:	Identified (Qvalue)

│ │ └─ Imputation Strategy:	Use Background Signal

│ ├─ Proteotypicity Filter:	None

│ ├─ Protein LFQ Method:	MaxLFQ

│ ├─ Quantity MS Level:	MS2

│ ├─ Quantity Type:	Area

│ ├─ Cross-Run Normalization:	True

│ │ ├─ Normalization Filter Type:	None

│ │ ├─ Normalization Strategy:	Local Normalization

│ │ └─ Row Selection:	Identified in at least 1 Run (Sparse)

│ ├─ Quantification window:	Not Synchronized (SN 17)

│ ├─ Interference Correction:	True

│ │ ├─ Only Identified Peptides:	True

│ │ ├─ Exclude All Multi-Channel Interferences:	True

│ │ ├─ MS1 Min:	2

│ │ └─ MS2 Min:	3

│ ├─ Major (Protein) Grouping:	by Protein Group Id

│ ├─ Minor (Peptide) Grouping:	by Stripped Sequence

│ ├─ Major Group Quantity:	Mean peptide quantity

│ ├─ Major Group Top N:	True

│ │ ├─ Max:	3

│ │ └─ Min:	2

│ ├─ Minor Group Quantity:	Sum precursor quantity

│ └─ Minor Group Top N:	False

├─ DIA Analysis\Workflow

│ ├─ Method Evaluation:	False

│ ├─ MS2 DeMultiplexing:	Automatic

│ ├─ Profiling Strategy:	iRT Profiling

│ │ ├─ Carry-over exact Peak Boundaries:	False

│ │ ├─ Profiling Row Selection:	Minimum Qvalue Row Selection

│ │ │ └─ Qvalue Threshold:	0.001

│ │ └─ Profiling Target Selection:	Profile only non-identified Precursors

│ │	├─ Identification Criterion:	Qvalue

│ │	└─ Threshold:	0.001

│ ├─ Run Limit for directDIA Library:	-1

│ └─ Unify Peptide Peaks Strategy:	Select corresponding Peak

├─ DIA Analysis\XIC Extraction

│ ├─ XIC IM Extraction Window:	Dynamic

│ │ └─ Correction Factor:	1

│ ├─ XIC RT Extraction Window:	Dynamic

│ │ └─ Correction Factor:	1

│ ├─ MS1 Mass Tolerance Strategy:	Dynamic

│ │ └─ Correction Factor:	1

│ └─ MS2 Mass Tolerance Strategy:	Dynamic

│	└─ Correction Factor:	1

├─ Pulsar Search\Identification

│ ├─ PSM FDR:	0.01

│ ├─ Peptide FDR:	0.01

│ ├─ Protein Group FDR:	0.01

│ ├─ directDIA Workflow:	directDIA+ (Deep)

│ └─ PTM Localization Filter:	False

├─ Pulsar Search\Labeling

│ └─ Channels:

│	├─ Channel 1:	False

│	├─ Channel 2:	False

│	└─ Channel 3:	False

├─ Pulsar Search\Modifications

│ ├─ Max Variable Modifications:	5

│ └─ Select Modifications:

│	├─ Fixed Modifications::

│	└─ Variable Modifications: :	Oxidation (M)

├─ Pulsar Search\Peptides

│ ├─ Enzymes / Cleavage Rules:	Trypsin/P

│ ├─ Digest Type:	Specific

│ ├─ Max Peptide Length:	52

│ ├─ Min Peptide Length:	7

│ ├─ Missed Cleavages:	0

│ └─ Toggle N-terminal M:	True

├─ Pulsar Search\Result Filters

│ ├─ Fragment Ions:

│ │ ├─ Ion AA Length:	True

│ │ │ └─ N:	3

│ │ ├─ Ion Charge:	False

│ │ ├─ Ion Loss Type:	False

│ │ ├─ Ion Type:	False

│ │ ├─ m/z :	True

│ │ │ ├─ Max:	1800

│ │ │ └─ Min:	300

│ │ └─ Relative Intensity:	True

│ │	└─ Min:	5

│ └─ Precursors:

│	├─ Amino Acids:	False

│	├─ Best N Fragments per Peptide:	True

│	│ ├─ Max:	10

│	│ └─ Min:	6

│	├─ Best N Peptides per Protein Group:	False

│	├─ Channel Count:	False

│	├─ FASTA Matched:	False

│	├─ Missed Cleavage:	False

│	├─ Modifications:	None

│	├─ Peptide Charge:	False

│	└─ Proteotypicity:	False

├─ Pulsar Search\Speed-Up

│ ├─ IM DFD Processing:

│ │ └─ Use Dynamic IM Peak Filter:	True

│ │	└─ Target TIC Fraction:	0.9

│ └─ MS2 Index:	Automatic

├─ Pulsar Search\Tolerances

│ └─ Tolerance Parameters:

│	├─ Thermo IonTrap:

│	│ ├─ Calibration Search:	Dynamic

│	│ │ ├─ MS1 Correction Factor:	1

│	│ │ └─ MS2 Correction Factor:	1

│	│ └─ Main Search:	Dynamic

│	│	├─ MS1 Correction Factor:	1

│	│	└─ MS2 Correction Factor:	1

│	├─ Thermo Orbitrap:

│	│ ├─ Calibration Search:	Dynamic

│	│ │ ├─ MS1 Correction Factor:	1

│	│ │ └─ MS2 Correction Factor:	1

│	│ └─ Main Search:	Dynamic

│	│	├─ MS1 Correction Factor:	1

│	│	└─ MS2 Correction Factor:	1

│	└─ TOF:

│	├─ Calibration Search:	Dynamic

│	│ ├─ MS1 Correction Factor:	1

│	│ └─ MS2 Correction Factor:	1

│	└─ Main Search:	Dynamic

│	├─ MS1 Correction Factor:	1

│	└─ MS2 Correction Factor:	1

└─ Pulsar Search\Workflow

├─ Fragment Ion Selection Strategy:	Intensity Based

├─ In-Silico Generate Missing Channels:	False

└─ Use DNN Predicted Ion Mobility:	Auto

END-SETTINGS

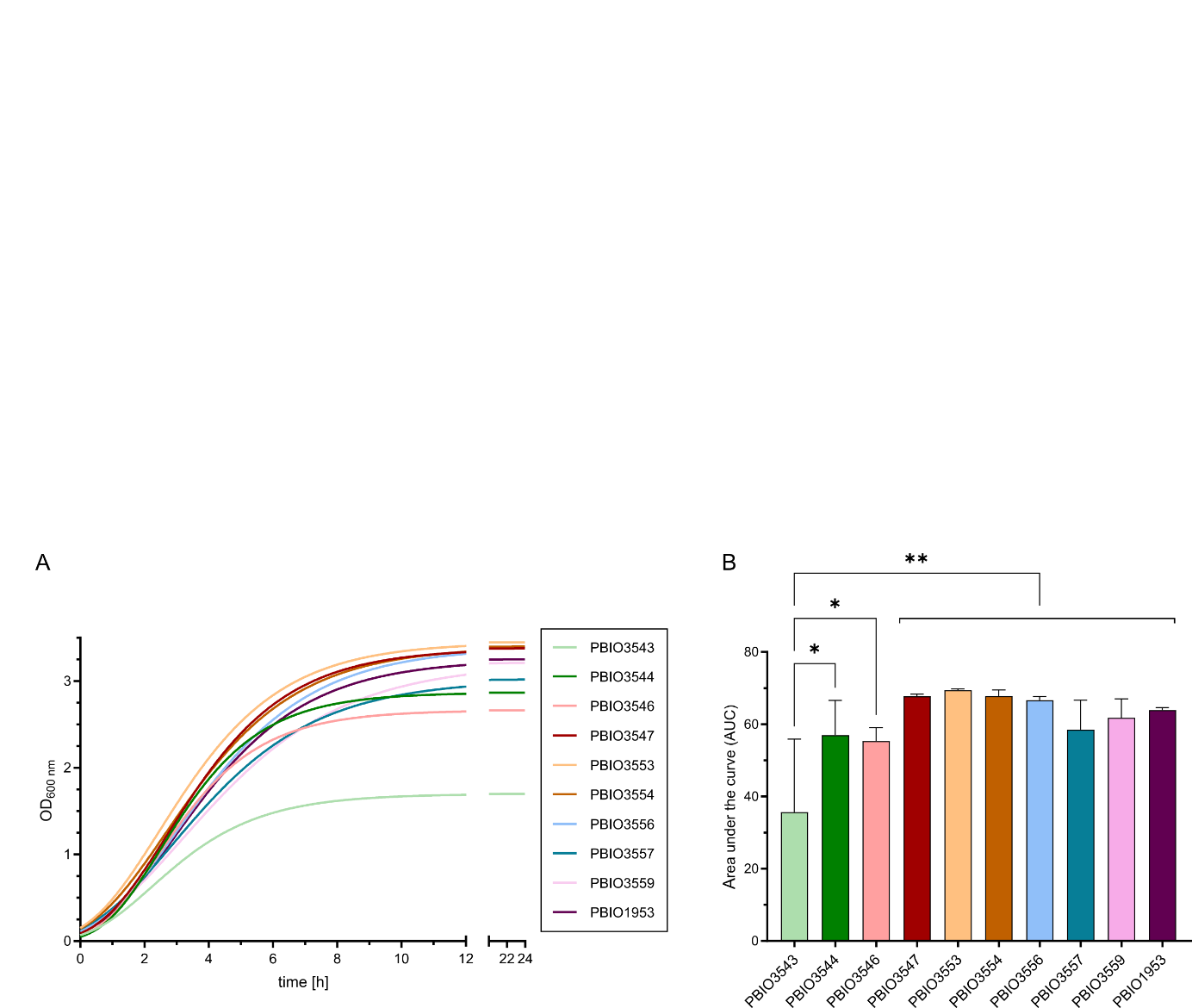

Figure S1: Growth kinetics of KP and KV strains in LB medium. Strains were cultivated in LB medium in biological triplicates. (A) Growth kinetics are displayed using GraphPad Prism and Gompertz-Growth fitting. (B) The area under the curve was calculated from three individual cultivations. The significance of differences was calculated using one-way ANOVA (p-value: * 0.33; ** 0.02).

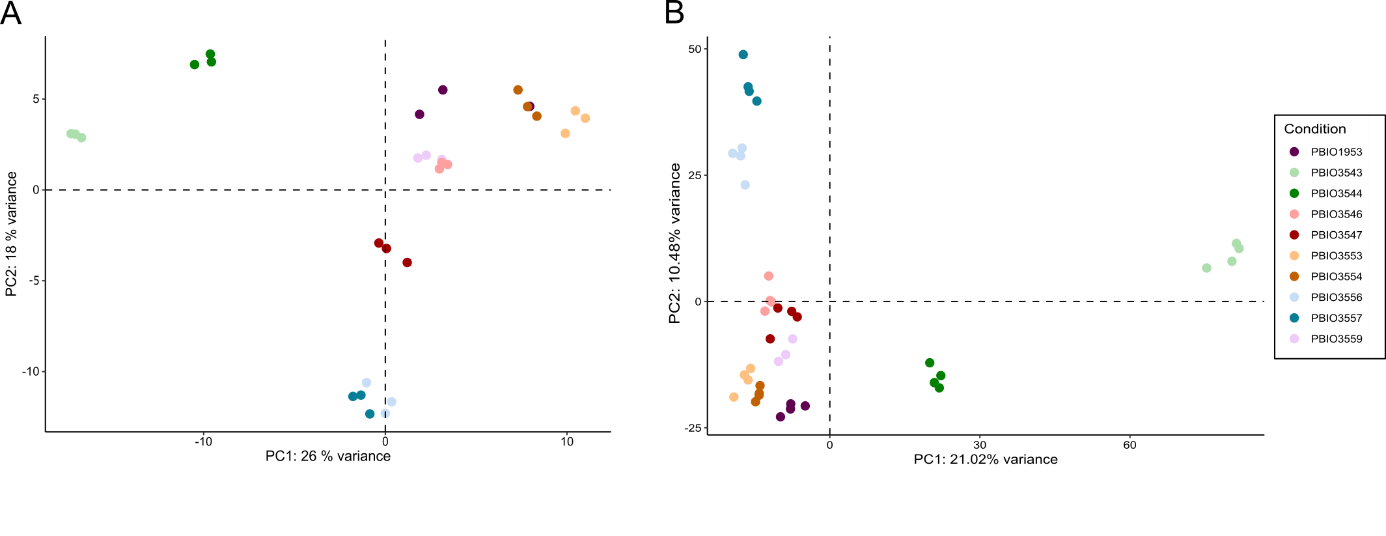

Figure S2: Principle component analysis. (A) Transcriptomic samples were separated by the PC1 (26% variance) and PC2 (18% variance). Data from three biological replicates are represented. (B) Protemomic samples were separated by the PC1 (21.02% variance) and PC2 (10.48% variance). Data from minimum three biological replicates are represented.

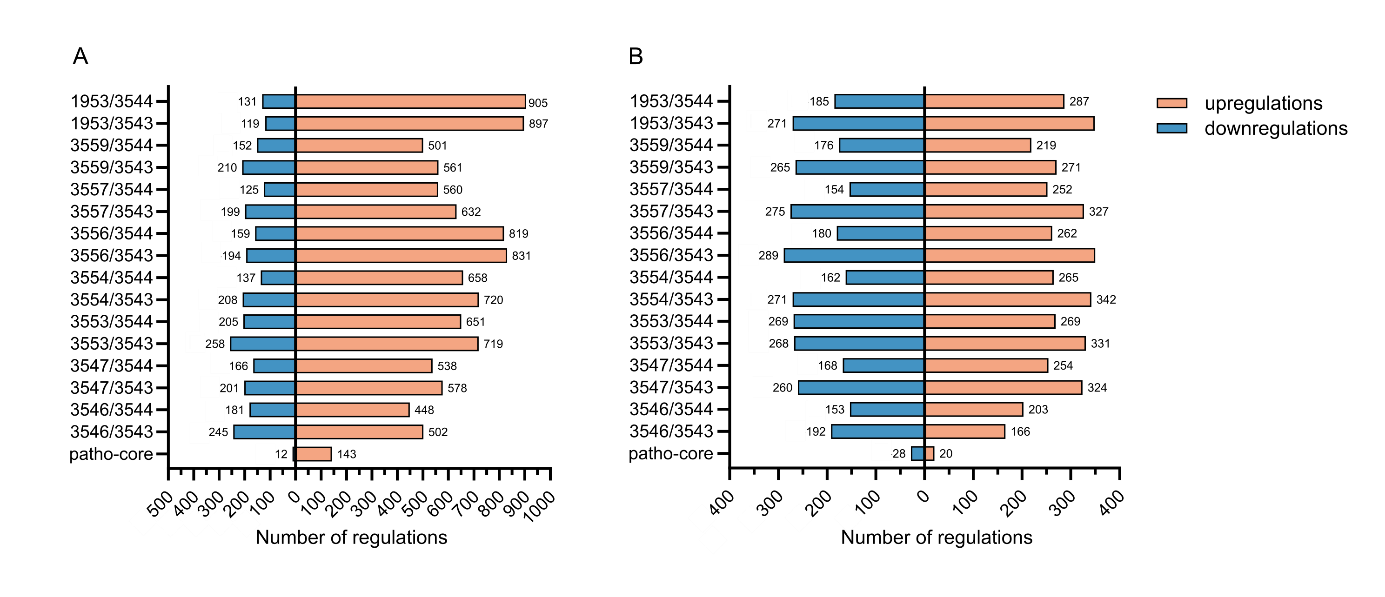

Figure S3: Number of strain specific and shared (patho-core) regulations on transcriptomic and proteomic levels. Differentially expressed genes were analyzed by comparing each pathogenic KP strain with each KV strain. Genes were assumed as differentially expressed when the comparison revealed a |L2FC|>1.5 and a significant p‑adjusted value of <0.05. Depending on the L2FC, genes were differentiated whether they are downregulated (L2FC<‑1.5) or upregulated (L2FC>1.5). The number of regulated genes on (A) transcriptomic and (B) proteomic levels is shown for each comparison and additionally the number of genes within the patho-core is indicated.

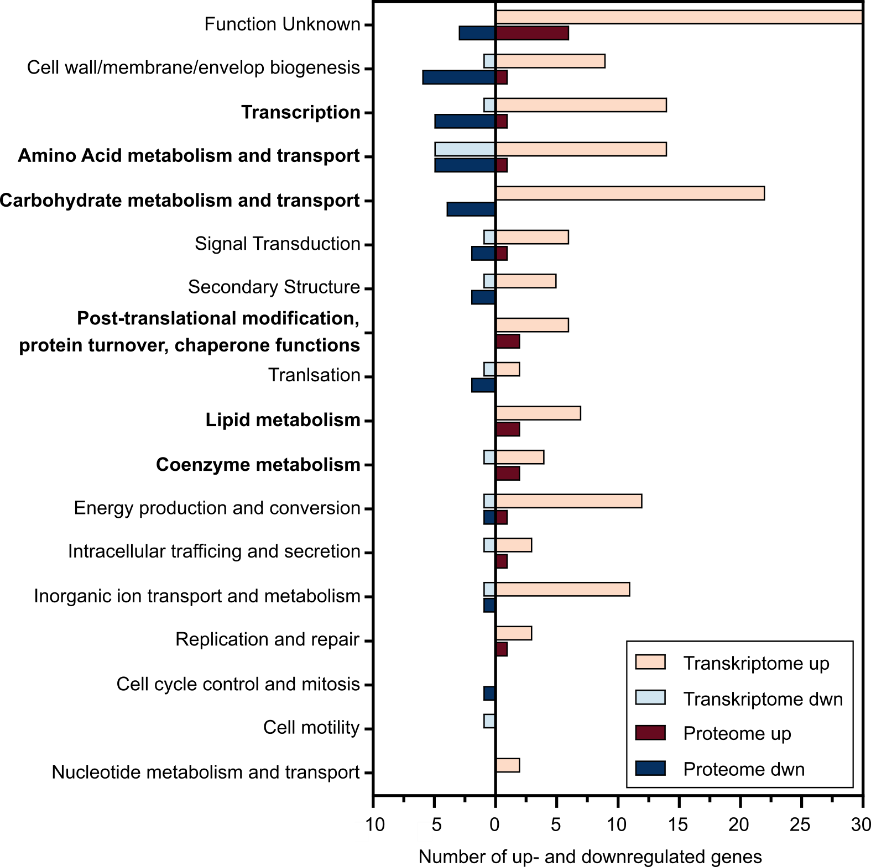

Figure S4: Number of up- and downregulations on transcriptomic and proteomic levels assigned to the different COG categories. Whenever available, information about the COG for all regulated genes on transcriptomic and proteomic levels were annotated based on the Bakta and EggNOG database and plotted separately on transcriptomic and proteomic levels and up- or down-regulated genes as indicated.

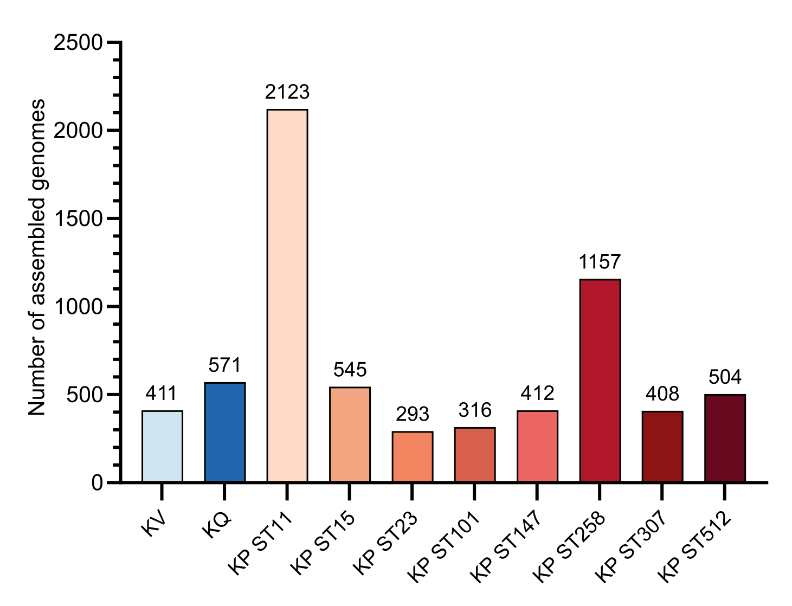

Figure S5: **Number of publicly available genomes used for the construction of a patho-core genome of KP.** The number of genomes downloaded from the NCBI database is shown for KV, KQ and each of the eight successful international high-risk KP STs.

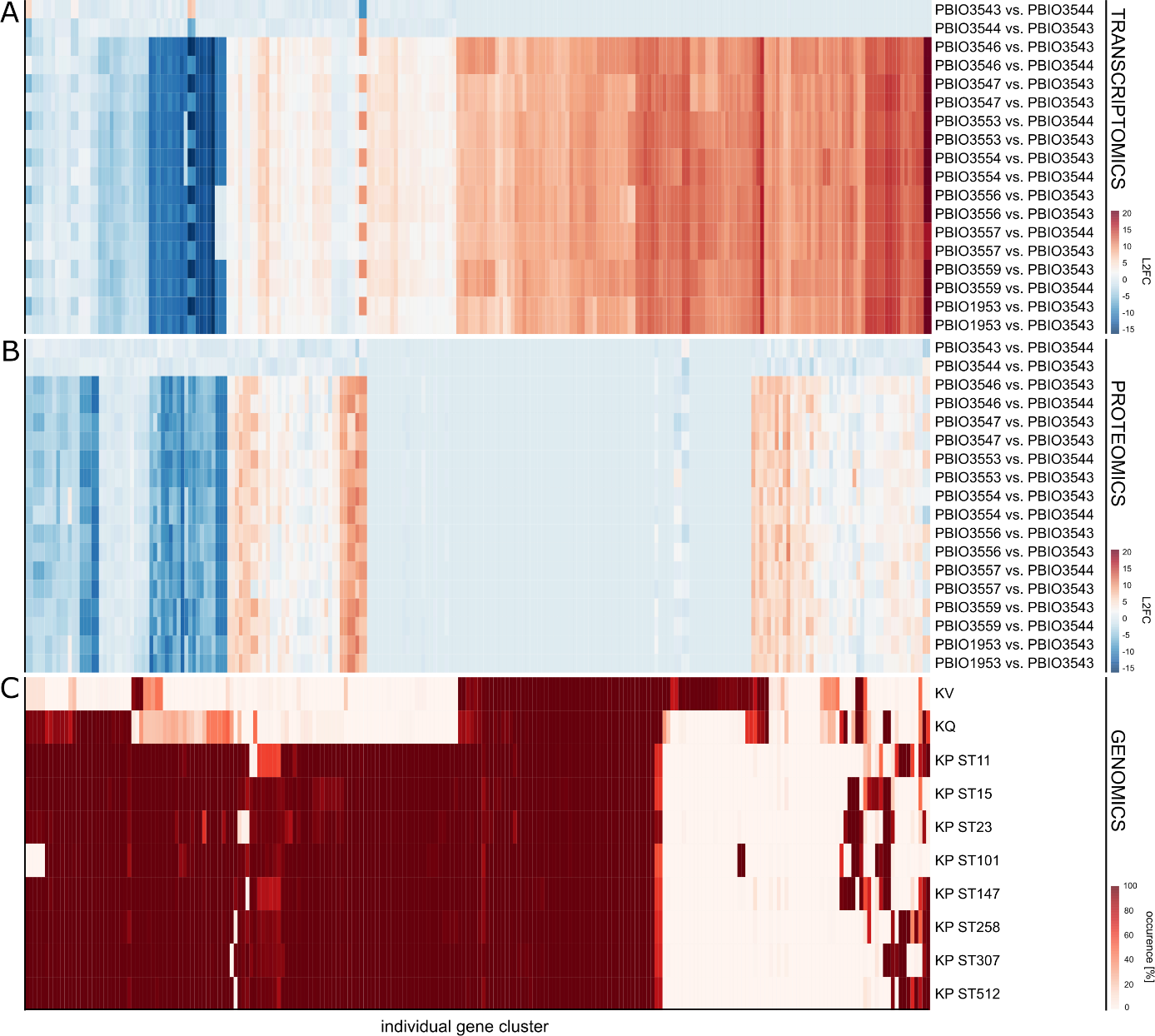

Figure S6: **Distribution of regulated genes within the large *Klebsiella* spp. pangenome.** Genes regulated either on transcriptomic and/or proteomic levels were mapped against the large pangenome of *Klebsiella* spp. including *K. variicola* (KV) and *K. quasipneumoniae* (KQ) and eight different KP STs (ST11, ST15, ST23, ST101, ST147, ST258, ST307 and ST512). The regulation on transcriptomic (A) and proteomic (B) levels is shown as the L2FC of the indicted comparisons each. (c) The occurrence of the individual gene clusters within the different *Klebsiella* species and STs of the large pangenome is shown in [%].
